# ETS-guided iPSC-endothelial models recapitulate malaria pathogenesis

**DOI:** 10.1101/2025.07.01.662615

**Authors:** François Korbmacher, Hannah Fleckenstein, Rory Long, Patryk Poliński, Livia Piatti, Borja López-Gutiérrez, Alina Batzilla, Dennis Crusius, Vikas Trivedi, Miki Ebisuya, Maria Bernabeu

## Abstract

The sequestration of the malaria parasite *Plasmodium falciparum* in the microvasculature is a major driver of severe malaria, but its pathogenic mechanisms still remain unknown. Advancements in induced pluripotent stem cell (iPSC) technologies offer unique opportunities to study parasite interactions with blood vessels in a well-defined host environment. However, endothelial iPSC-differentiation methods often result in cells with mixed epithelial identity. Here, we have generated an iPSC line with inducible and simultaneous expression of ETS transcription factors (ETV2, FLI1, ERG), which resulted in improved endothelial cell identity and strong barrier function. These cells display a high affinity to infected red blood cells. Exposure to parasite products caused significant endothelial metabolic changes and splicing alterations. Furthermore, it disrupted the iPSC-endothelial barrier, as a consequence of transcriptional downregulation of key barrier processes, and alteration of severe malaria biomarkers. Our novel iPSC-based approach represents a new *in vitro* platform to study the pathogenesis of vascular infections.

## Introduction

*Plasmodium falciparum* is the most severe human malaria parasite, causing over 600,000 deaths annually (*1*). Severe malaria affects multiple organs, including the lungs, kidneys, liver, heart, and brain, with cerebral malaria being one of the most fatal complications. A major contributor to organ pathogenesis is *P. falciparum*-infected red blood cell (infected RBC) accumulation in the microvasculature, a phenomenon also known as sequestration. Models of severe malaria comprise either *in vitro* or rodent malaria infection models, with the latter poorly recapitulating high levels of parasite sequestration reported in patients. Since the endothelium is the key interface for pathogenic interactions, most *in vitro* experimental approaches have focused on endothelial cells. Unfortunately, immortalised cell lines often lack crucial junctional markers, while primary cells are prone to rapid dedifferentiation and batch-to-batch variability (*2*). Moreover, both choices present poor endothelial barrier function, a critical feature for understanding vascular pathogenesis in severe malaria.

Induced pluripotent stem cells (iPSCs) have become a promising alternative to address these shortcomings. Differentiation methods have been widely used in malaria research to generate RBC from hematopoietic stem cells (*3, 4*). More recently, iPSC-derived erythrocytes have emerged as a novel tool for studying *P. falciparum* invasion into RBC (*5*), a technology that could be extended to dissect host-parasite interactions with blood vessels. One of the most commonly used endothelial differentiation methods generates iPSC-derived brain microvascular endothelial cell-*like* cells (iBMEC-*like*). This differentiation method has been popular, as it is easy to implement and yields cells with consistent and strong barrier properties, functionally mimicking those found at the blood-brain barrier (BBB) (*6, 7*). However, iBMEC-*like* cells have recently sparked controversy, particularly with regard to their endothelial identity, as they express epithelial markers (*8*). Nevertheless, iBMEC-*like* cells continue to be used in various applications (*9–11*), including the first iPSC *in vitro* model of cerebral malaria (*12*).

Lentiviral expression of ETS transcription factors has been proposed as a strategy to rescue the endothelial phenotype of iBMEC-*like* cells (*8*), and their overexpression is recently becoming a popular methodology for iPSC differentiation (*13–17*). Among ETS transcription factors, ETV2, ERG, and FLI1 are regulators of endothelial development. ETV2 stands out as a key orchestrator, coordinating the development by activating specific endothelial gene expression and even enabling the direct conversion of somatic cells into endothelial cells (*8, 18, 19*). ERG and FLI1 are both described as essential factors in endothelial development, angiogenesis, and maintenance (*20, 21*). However, to date, differentiation systems that promote ETS transcription factor overexpression have not been applied to investigate malaria pathogenesis, neither via lentiviral transduction nor through the introduction of stable transgenes. Here, we generated a doxycycline-inducible, stable transgenic iPSC line that upregulates ETV2, ERG, and FLI1 during iBMEC endothelial differentiation. The resulting cell type, ETS-iBMEC, presented increased transcription of endothelial markers and downregulation of epithelial transcripts. Although ETS-iBMECs showed equally strong barrier properties as iBMEC-*like* cells when grown in a 3D model, they exhibited a marked shift towards endothelial ultrastructural morphology. Perfusion of *P. falciparum*-infected RBCs displayed increased cytoadhesion to ETS-iBMECs, and parasite products induced endothelial barrier disruption, two key pathogenic hallmarks of severe and cerebral malaria. Moreover, incubation with parasite egress product induced a progressive transcriptional dysregulation of endothelial development genes, robust downregulation of junctional, cytoskeletal, and basement membrane anchor genes in ETS-iBMECs. Interestingly, we also observed a dysregulation of spliceosome and metabolic pathways, revealing potential additional mechanisms for endothelial disruption.

## Results

### ETS transcription factors promote endothelial differentiation in a DOX-inducible iPSC line

We developed an endothelial differentiation protocol based on the widely used iBMEC-*like* protocol, including the addition of hypoxia, which has been shown to improve barrier functionality (Fig. 1) (*7, 22*). To boost the endothelial identity of iBMEC-*like* cells, we genetically engineered the iPSC(IMR90-4) line with a doxycycline (DOX)-inducible Tet-One system to overexpress ETV2, ERG, and FLI1 (ETS) transcription factors (Fig. 1A). This was achieved through a piggyBac transposase, which facilitated genomic integration of the three inducible ETS factors (Fig. S1A). A clone exhibiting robust induction of all three factors was selected (Fig. S1B). Differentiation of the DOX-induced cells (hereafter referred to as ETS-iBMECs) was performed using the standard iBMEC-*like* protocol, with DOX-induction of ETV2, ERG, and FLI1 from day 6 to day 11 (Fig. 1B), during the endothelial expansion phase (*7, 8*). To rule out silencing of the Tet-One system over the course of differentiation, we generated a parallel DOX-inducible iPSC line expressing fluorescent reporters. Fluorescence imaging confirmed stable signal intensity throughout the differentiation timeline, validating the robustness of the DOX-induction system (Fig. S1C).

**Fig. 1.**
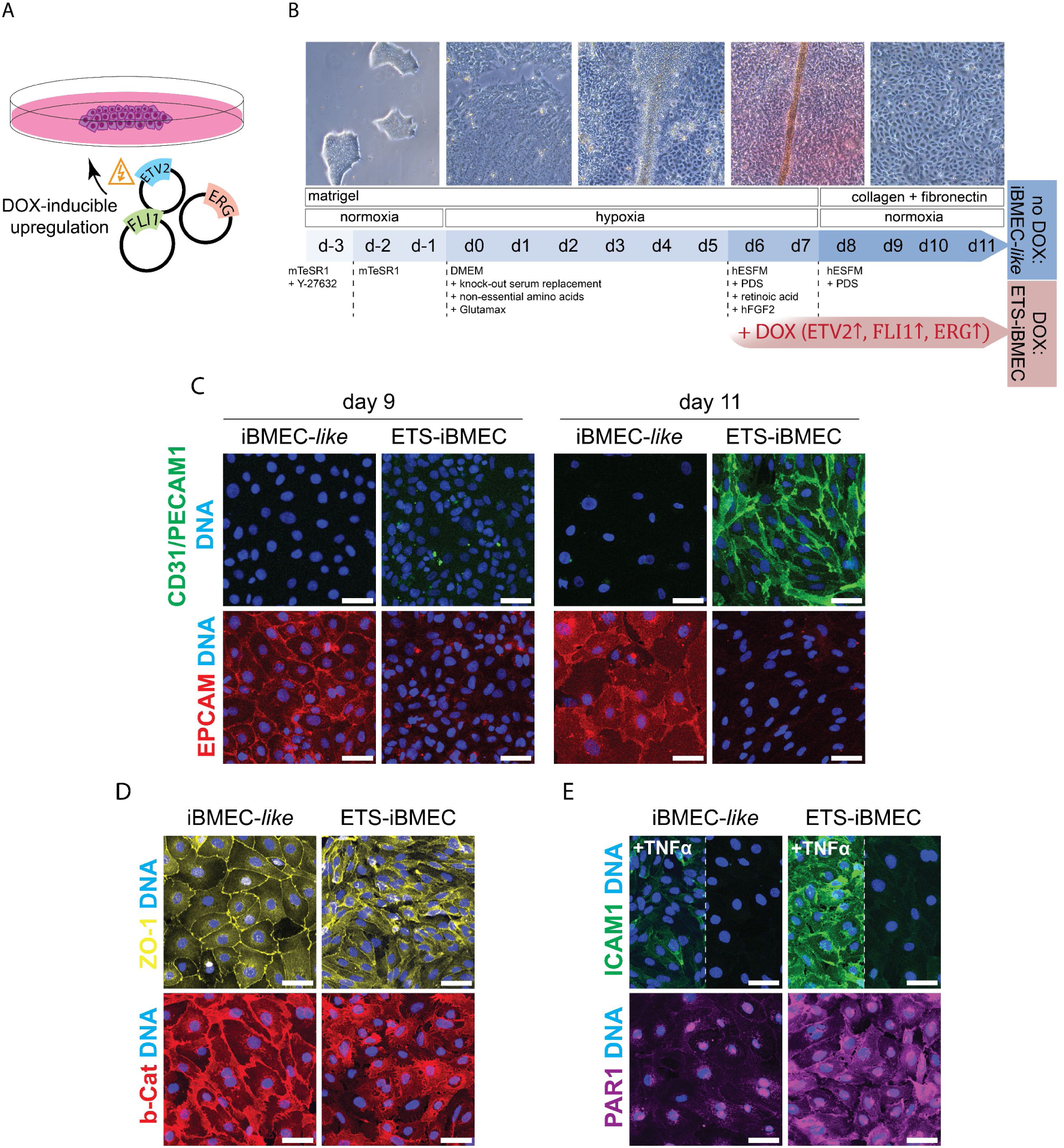
Generation of the ETS-iBMECs. (**A**) The stable transgenic line was generated by transfecting iPSCs with a DOX-inducible upregulation system for the ETS-transcription factors: ETV2, ERG, and FLI1. (**B**) During the iBMEC-*like* differentiation protocol, the ETS-transcription factors were DOX-induced from day 6 until day 11 to generate ETS-iBMECs. A schematic representation of the iBMEC-*like* differentiation is shown above. (**C**) Immunofluorescence confocal microscopy of ETS-iBMEC and iBMEC-*like* cells stained with antibodies against epithelial EPCAM (red), endothelial CD31/PECAM1 (green), and nuclei (blue; DAPI) in samples fixed at day 9 and day 11 after differentiation. (**D**) Immunofluorescence confocal imaging of ETS-iBMEC and iBMEC-*like* cells with antibodies against tight junction protein ZO-1 (yellow), and adherens junction protein β-catenin (red). (**E**) Immunofluorescence staining of the endothelial receptor ICAM1 (green), with and without prior TNFα stimulation, and the thrombin receptor PAR1 (purple) and nuclei (blue, DAPI). Scale bars, 100 µm.

To fully assess the robustness of the ETS-driven differentiation, we characterised the endothelial identity of the DOX-induced cells (ETS-iBMECs) and benchmarked it against the non-induced control (iBMEC-*like*) (Fig. 1B) using immunofluorescence, transcriptomics, barrier functional assays, and electron microscopy. Immunofluorescence assays showed a consistent loss of epithelial EPCAM over the course of DOX induction, being absent at day 11. Conversely, the expression of the endothelial marker CD31 was significantly increased at day 11 in ETS-iBMECs (Fig. 1C). Furthermore, labelling of the tight junction marker ZO-1 and adherens junction protein β-Catenin revealed notable changes in cell morphology, with DOX-induced cells becoming smaller and elongated after ETS induction (Fig. 1D). Moreover, the ETS-induced cells responded to TNFα endothelial activation by increasing the expression of ICAM1, a key endothelial receptor responsible for recruitment of *P. falciparum-*infected RBCs and leukocytes (*23*). Similarly, the thrombin receptor PAR1, associated with severe malaria through dysregulation of EPCR (*24*), displayed increased expression following DOX-induction (Fig. 1E). Together, these results show that the ETS-induced stable iPSC line promotes a shift towards endothelial identity, while non-induced iBMEC-*like* cells retain features reminiscent of epithelial cells.

### ETS-iBMECs present increased expression of endothelial transcripts

To dissect the transcriptional changes triggered by ETV2, ERG and FLI1 upregulation, we performed bulk RNA sequencing (RNA-seq) and compared the expression profiles of ETS-iBMECs with their non-induced counterpart and primary brain microvascular endothelial cells (primary brain ECs). Differential gene expression analysis confirmed the overexpression of ETV2, ERG, and FLI1, and revealed that ETS-iBMECs upregulate genes associated with endothelial identity (*CDH5, PECAM1, FLT1, FLT4, TIE1*), as well as genes involved in angiogenesis and vascular development (*ANGPT2, TEK, ENG, APLN, ROBO4, EGFL7, SCUBE1, VASH1, RASIP1*), together with the vascular tight junction transcript *CLDN5*. Conversely, many epithelial genes were found to be downregulated (*CDH3, ELF3, EPCAM, ESRP1, GRHL2, OVOL1, OVOL2, TP63*) (Fig. S2), highlighting a clear transcriptional divergence between ETS-iBMEC and iBMEC-*like* cells. Gene ontology (GO) over-representation analysis further supports the shift towards an improved endothelial identity, highlighting biological processes involved in angiogenesis, endothelial proliferation, blood circulation, and cell adhesion being upregulated in ETS-iBMEC (Table S1). Indeed, the top five most significantly upregulated biological terms include angiogenic, circulatory, and vascular developmental processes, reflecting a strong shift toward endothelial programming after ETS induction (Fig. 2A). Additional upregulated clusters included pathways involved in signal transduction and cellular homeostasis, such as G-protein coupled pathways, ion transport, and tissue homeostasis, alongside cell physiological processes related to chemotaxis and migration, cell adhesion, and synaptic organization (Fig. 2B). Notably, the synaptic organisation cluster includes several genes expressed by endothelial cells, such as *NRP1*, *NRP2*, *PLXND1*, *RELN*, and *NLGN2,* that are key regulators of angiogenesis and neurovascular interactions (Table S1). DOX induction also enhanced the expression of genes involved in immune activation, highlighting the potential of ETS-iBMECs as an *in vitro* platform for modelling inflammatory responses, a relevant feature for infection models. Conversely, mainly non-endothelial morphogenetic processes were downregulated, including meso- and ectodermal developmental processes of epidermis, skin, glands, bones and muscle tissue as the most significant decreased processes (Fig. 2C). Among these, epithelial developmental processes were found to be downregulated, together with terms associated with cell-contact organisation, extracellular organisation, cell motility, and multicellular organisation (Fig. 2D).

**Fig. 2.**
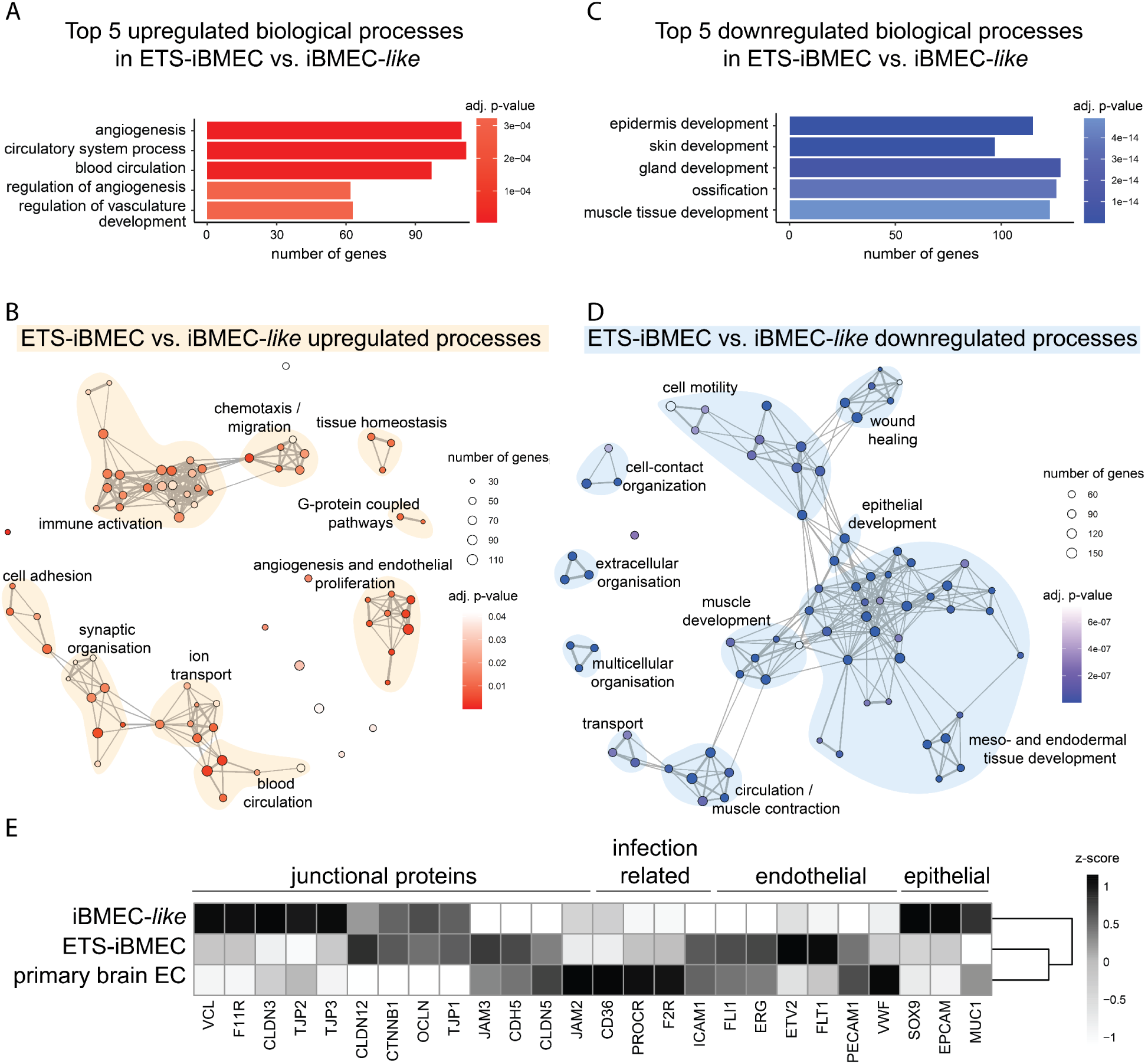
Transcriptional analysis of bulk RNA-seq in ETS-iBMECs (DOX), iBMEC-*like* (no DOX) and primary brain ECs. (**A**) Top 5 significantly upregulated GO biological processes and (**B**) GO term over-representation analysis map of differentially upregulated genes on ETS-iBMEC compared with iBMEC-*like*. (**C**) Top 5 significantly downregulated GO biological processes and (**D**) GO term over-representation analysis map for differentially downregulated genes on ETS-iBMEC compared with iBMEC-*like*. The GO term maps show the 75 most significant GO term over-representation analyses from biological processes, which include ≥ 30 differentially expressed genes with adjusted p-value < 0.05 and log_2_FC > ±1. Node connections indicate shared genes, and node clusters are manually annotated. (**E**) Normalised gene expression of a selection of endothelial and epithelial genes across iBMEC-*like* cells, ETS-iBMEC and primary brain endothelial cells (primary brain EC). Z-score scaled by column (gene).

To further compare the expression profiles of the induced cell line with primary brain ECs, a widely used model for malaria studies, we analysed a curated panel of key endothelial and epithelial genes, along with markers important for modelling severe malaria pathogenesis (Fig. 2E). This analysis revealed a similar expression of these markers in ETS-iBMECs and primary brain ECs compared to iBMEC-*like* cells. ETS-iBMECs and primary brain ECs both exhibited similar expression levels of the endothelial marker *PECAM-1* (CD31) and did not express epithelial genes such as *MUC1*, *EPCAM* and *SOX9*. Endothelial transcription factors *ERG* and *FLI1* reached comparable levels of expression in both primary brain ECs and ETS-iBMECs. However, ETV2 was expressed at higher levels in DOX-induced cells compared to primary brain ECs, an expected phenotype due to the downregulation of ETV2 in fully differentiated endothelial cells. Analysis of junctional proteins showed that DOX-induced ETS-iBMECs partially recapitulate the expression profile of primary brain ECs. Junctional genes like *CLDN5* (Claudin-5), *CDH5* (VE-Cadherin), and *JAM3* showed similar expression in both ETS-iBMECs and primary brain ECs, while other important endothelial tight junctional genes like *TJP1* (ZO-1), and *OCLN* (Occludin) showed a higher expression in ETS-iBMECs, suggesting an improvement in endothelial barrier properties in DOX-induced cells. Lastly, parasite-interacting receptors such as *PROCR* (EPCR) and *ICAM1* show increased expression following DOX induction, except for *CD36*. Together, these transcriptional changes highlight a broad transcriptional shift in ETS-iBMECs, with an expression pattern that more closely resembles primary brain ECs (Fig. 2E).

### ETS-iBMEC microvessels lose epithelial ultrastructural features while maintaining barrier function

To assess the microvascular properties of ETS-iBMECs, we incorporated them into a previously established 3D microfluidic device consisting of a pre-patterned type I collagen hydrogel, generated through soft lithography and injection moulding (*25*). Primary human astrocytes (HA) and brain vascular pericytes (HBVPs) were seeded at a 7:3 ratio in the collagen bulk (*26*), while endothelial cells were seeded into the microfluidic network by gravity-driven perfusion (Fig. 3A). After a few days in culture, the endothelial cells formed perfusable microvessels with sparse contacts to the surrounding glial and perivascular cells. Both iBMEC-*like* cells and ETS-iBMECs formed perfusable microvessels in this setup (Fig. 3B). Nevertheless, seeding required a pre-coating with fibronectin and collagen IV and was more challenging than with primary brain ECs, resulting in much lower throughput. To determine barrier function, we measured the apparent trans-endothelial permeability following perfusion with 40 kDa and 10 kDa dextran (Fig. 3C). Remarkably, ETS-iBMEC microvessels reached permeability values of <10^-8^ cm/s with median levels of 1.5x10^-8^ cm/s, representing an over 100-fold barrier improvement over primary brain EC microvessels that present median permeability values of 4.3x10^-6^ cm/s in the same setup. Furthermore, both iBMEC-*like* and ETS-iBMEC microvessels were largely impermeable to smaller molecular weight tracers (Fig. 3D-E, Fig. S3A), with ETS-iBMECs showing slightly elevated 10 kDa dextran permeability rates (median 5.9x10^-8^ cm/s) compared to iBMEC*-like* controls (median 8.8x10^-9^ cm/s). These results demonstrate that microvessels made from iBMEC*-like* and ETS-iBMECs possess stronger barrier properties than those derived from primary brain ECs.

**Fig. 3.**
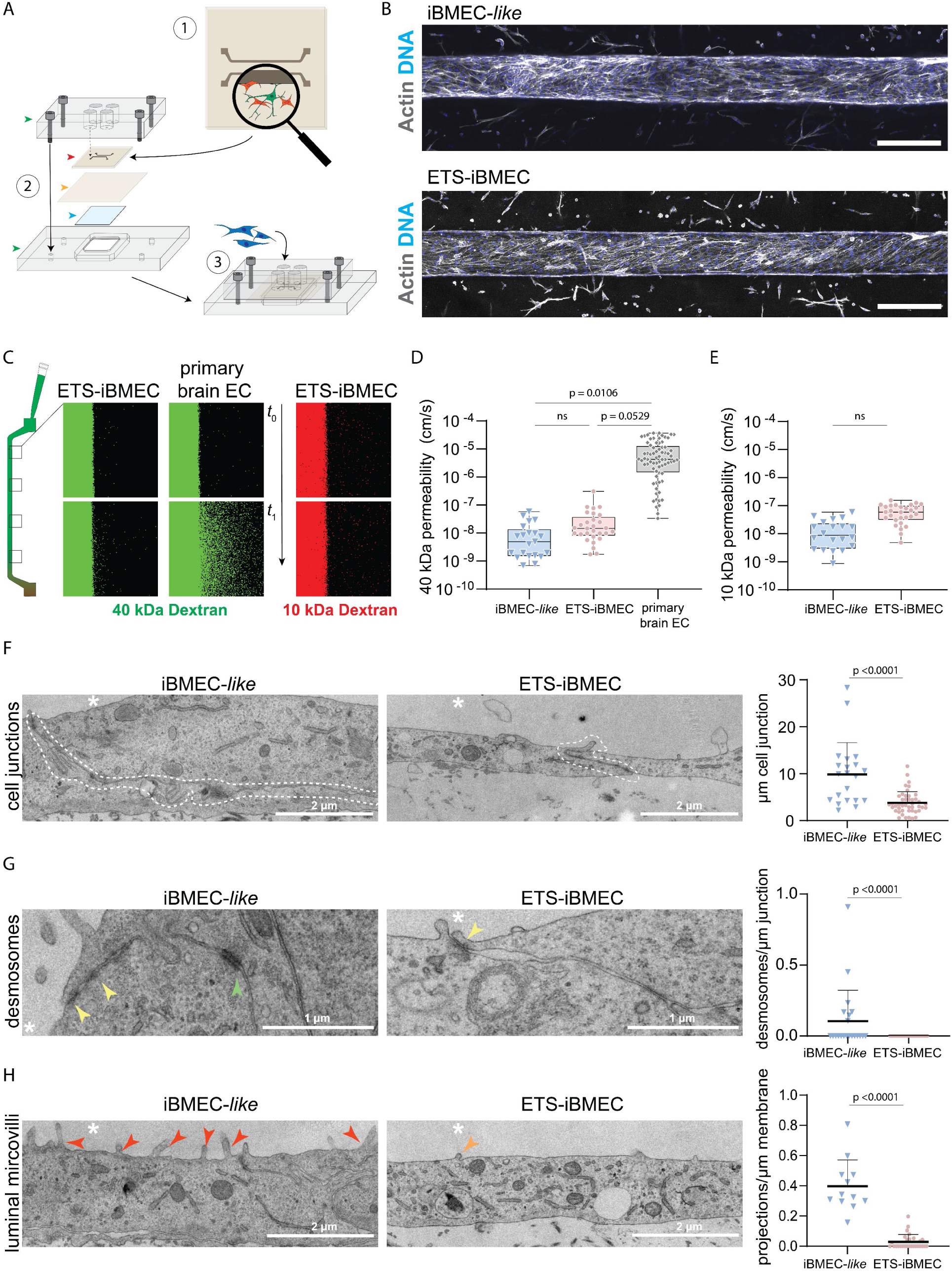
Barrier and ultrastructural morphological characterisation of ETS-iBMEC (DOX) and iBMEC-*like* cells (no DOX). (**A**) Schematic of a 3D-engineered microfluidic device composed of a collagen hydrogel containing primary astrocytes (green cells) and pericytes (red cells). The upper collagen layer (red arrow) includes two pre-patterned, parallel microchannels (1), and is assembled with a flat lower collagen layer (yellow) and a coverslip (blue) between two polymeric jigs (green) (2). The top jig has inlet and outlet wells, which are aligned to the entry and exit of each microchannel, allowing for direct perfusion. ETS-iBMECs, iBMEC-*like* and primary brain ECs are seeded along the microchannel by luminal gravity perfusion through the inlet (3). (**B**) The endothelial cells generate tubular microvessels. Top: iBMEC-*like* microvessels, bottom: ETS-iBMEC microvessels both stained for actin (phalloidin, grey) and DNA (DAPI, blue). Scale bar: 100 µm. (**C**) Representative images of barrier permeability assay on 3D microvessels perfused with fluorescently labelled 40 kDa FITC dextran (green) and 10 kDa AF647 dextran (red). Analysis was done in squared regions of interest (ROI) along iBMEC-*like*, ETS-iBMEC and primary brain EC microvessels at two different timepoints of perfusion (t_1_ = 150 seconds). (**D**, **E**) Quantified apparent permeability to 40 kDa dextran and 10 kDa dextran. Each dot represents individual ROIs in iBMEC-*like* (N = 3 microvessels), ETS-iBMECs (N = 4 microvessels) and primary brain ECs (N = 8 microvessels). Statistical analysis performed with Dunn’s multiple comparisons test. Boxplots indicate median and interquartile range (**F** to **H**). TEM images of iBMEC-*like* and ETS-iBMEC microvessels. Scale bars: 2 µm. (**F**) Representative images of junctions outlined with a white dotted line (left and center), and quantification of differences in junctional length (right). Statistical significance calculated by Mann-Whitney U test. iBMEC-*like* (N = 21 junctions), and ETS-iBMEC (N = 45 junctions). (**G**) Representative images of tight junctions (yellow arrows), characterised by tightly connected electron-dense membranes, and desmosomes with electron-dense plaque and wider intermembrane space (green arrow). Statistical significance of the presence of desmosomes calculated by Mann-Whitney U test. iBMEC-*like* (N = 21 junctions), ETS-iBMEC (N = 45 junctions). (**H**) Representative images of microvilli (red arrow) and smaller projections (orange arrow). Statistical difference in the quantification of all projections calculated by Mann-Whitney U test. iBMEC-*like* (N = 28 ROIs) and ETS-iBMEC (N = 12 ROIs).

While both differentiated cell types exhibit robust barriers, their transcriptional profiles are markedly distinct. To better understand these differences, we conducted transmission electron microscopy (TEM) on thin sections of the 3D brain microvascular model. IBMEC-*like* cells showed greater apical-basal height, forming elongated junctions characterized by the typical architecture of epithelial junctional complexes: apical tight junctions followed by adherens junctions, prominent desmosomes, and interdigitated regions (Fig. 3F, G). Conversely, ETS-iBMECs were flatter and had shorter junctions (Fig. 3F), containing tight junctions, and no desmosomes were observed (Fig. 3G). Additionally, the two cell types exhibited distinct apical surface morphologies: iBMEC-*like* cells displayed epithelial microvilli-like projections, which were rarely observed in ETS-iBMECs (Fig. 3H). The ultrastructural morphology of ETS-iBMECs closely resembled that of primary brain ECs (*27*) (Fig. S3B), further suggesting a loss of epithelial characteristics. Altogether, these findings demonstrate that ETS-iBMECs outperform primary brain ECs in barrier function, while displaying enhanced endothelial transcription and ultrastructure compared to iBMEC-*like* cells.

### *P. falciparum* presents strong binding to ETS-iBMECs under flow

Cytoadhesion of parasite*-*infected RBCs to endothelial EPCR and ICAM1 is a key determinant of severe and cerebral malaria (*28–32*). To evaluate the suitability of ETS-iBMECs for malaria *in vitro* pathogenesis studies, we first tested their capacity to bind *P. falciparum-*infected RBCs. Fluorescently labelled *P. falciparum-*infected RBCs were perfused for 30 minutes through microfluidic flow chambers seeded with either iBMEC-*like*, ETS-iBMEC, or primary brain ECs (Fig. 4A). Unbound parasites were washed out for 10 minutes and bound parasites were quantified after fixation. The *P. falciparum* line ITGICAM1 is known for its dual binding to CD36 and ICAM1 (Fig. S4, 4B), and showed significantly higher adhesion to ETS-iBMECs compared to iBMEC-*like* cells, and a non-significant increase in binding relative to primary brain ECs (Fig. 4C-D). The IT4var19 parasite line, known for strong affinity to endothelial EPCR (Fig. S4, 4B), a phenotype highly enriched in severe malaria patients, exhibited robust cytoadhesion to all three cell types (Fig. 4C-E). Notably, IT4var19 showed increased binding to ETS-iBMECs compared to iBMEC-*like* cells and primary brain ECs, although the difference was not statistically significant (Fig. 4E). Overall, the improved endothelial identity and the strong binding rates for *P. falciparum-*infected red blood cells exemplify the usefulness of ETS- iBMECs as an alternative cell model for malaria pathogenesis.

**Fig. 4.**
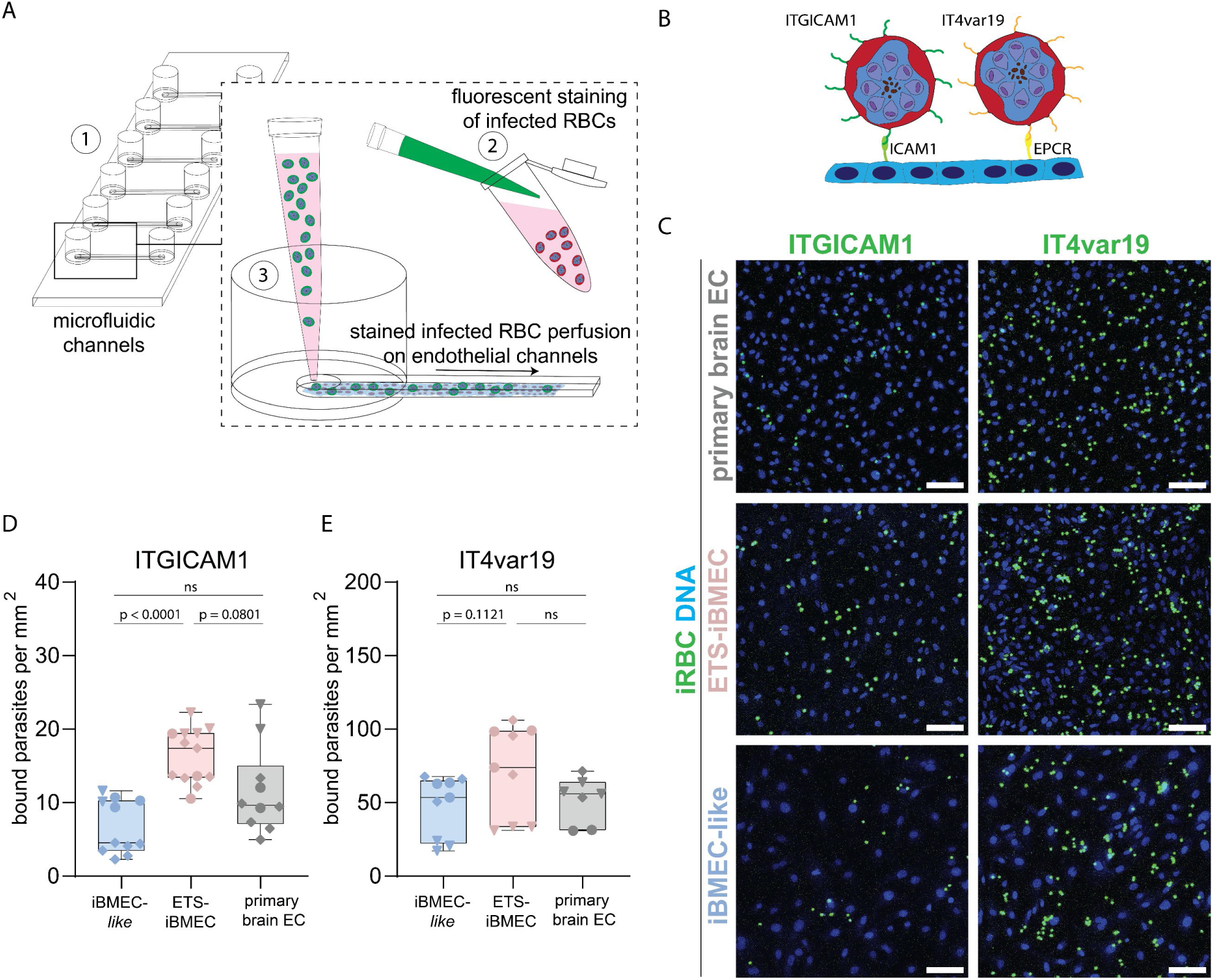
*P. falciparum in vitro* infection modelling for parasite cytoadhesion. (**A**) Endothelial channels are formed in microfluidic slides previously seeded with iBMEC-*like*, ETS-iBMEC and primary brain EC cells (1). Infected red blood cells (infected RBCs) are purified and fluorescently stained (2) and gravity-perfused through the endothelial channels, where they adhere under flow to endothelial cells (3). (**B**) PfEMP1 variants ITGICAM1 binds to the endothelial surface receptor ICAM1 and IT4var19 to EPCR. (**C**) Representative images of parasite cytoadhesion assay to iBMEC-*like*, ETS-iBMEC and primary brain EC channels. Infected RBCs in green, endothelial nuclear stain in blue. Scale bar: 100 µm. (**D**, **E**) Quantification of bound infected RBCs for *P. falciparum* lines ITGICAM1 (**D**) and IT4var19 (**E**) in microfluidic iBMEC-*like*, ETS-iBMEC and primary brain EC channels. Dot plot shows *P. falciparum-*infected RBC average binding to regions of interest, normalised per area (mm^2^), of three independent experiments including different differentiations and independent parasite purifications. Boxplots indicate the median and interquartile range. Statistical analysis performed with Dunn’s multiple comparisons test. Each dot corresponds to averaged binding across one microchannel.

### *P. falciparum* induces transcriptional changes in ETS-iBMECs, including metabolic alterations and alternative splicing events

Next, we aimed to identify endothelial disruptive mechanisms upon exposure to *P. falciparum-*infected RBCs. Accumulation of infected RBCs in the microvasculature results in the localised release of parasite-derived products near the endothelium at the late stages (schizonts) of the blood stage cycle, an event previously characterised as highly disruptive to barrier integrity in primary endothelial cells of various tissues (*26, 27, 33–35*). Here, we generated a solution containing parasite egress products by arresting the egress of highly synchronised late-stage *P. falciparum*-infected RBCs using the reversible protein kinase G inhibitor, compound 2 (C2), followed by the simultaneous release of parasite-derived products upon its removal (Fig. 5A). To understand the molecular mechanisms upon exposure to *P. falciparum-*iRBC products, we performed bulk RNA-seq after 8 and 24 hours of exposure to 5×10⁷ egressed infected RBCs/ml. As a control, we also sequenced iBMEC-*like* and primary brain ECs under the same conditions. While most of the dysregulated transcripts were unique to each cell type, some commonalities were observed between all three or in pairwise comparisons, suggesting that these pathways are commonly disrupted by the malaria egress products (Fig. S5A-B).

**Fig. 5.**
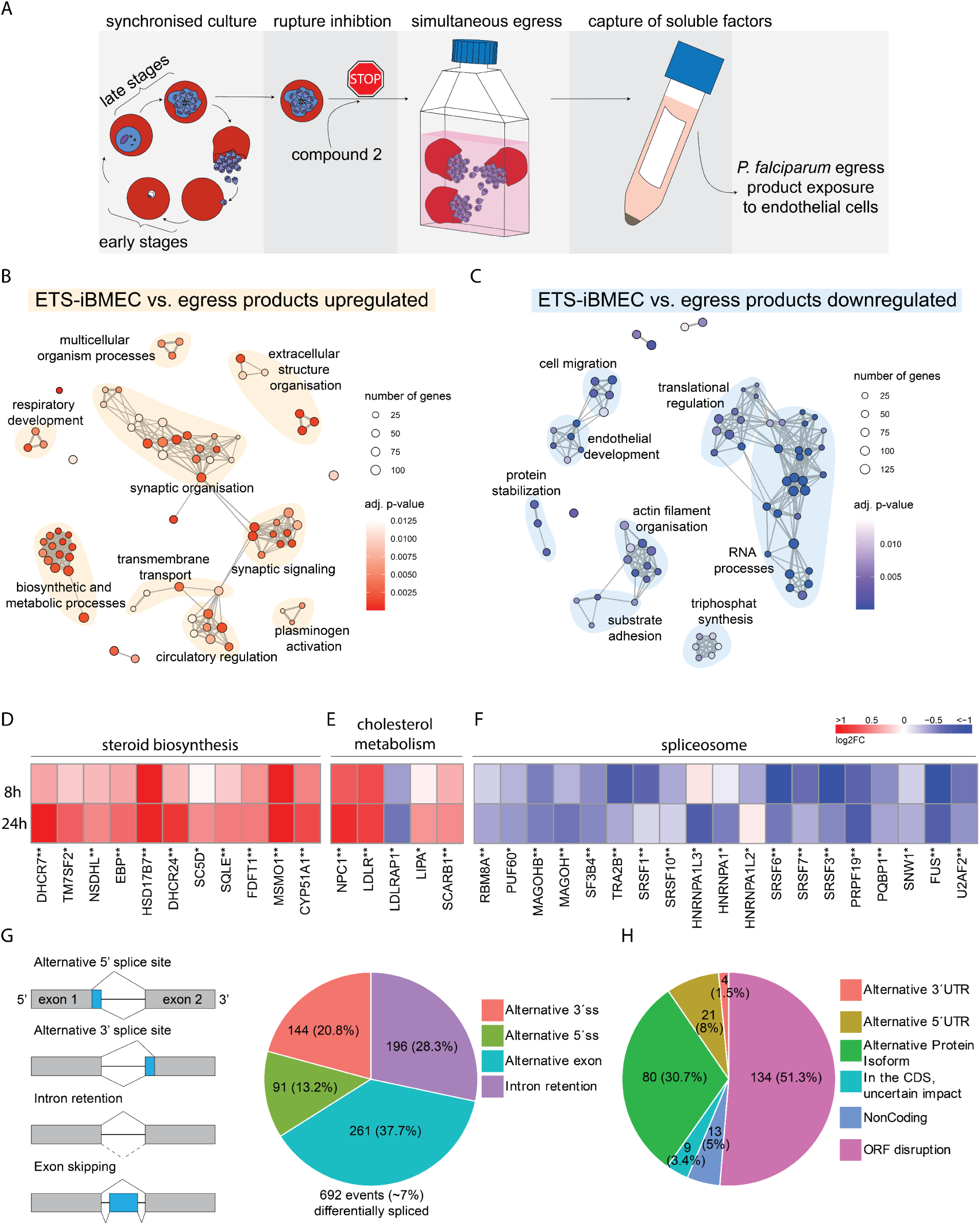
Transcriptional profiling of ETS-iBMEC response to malaria egress product exposure. (**A**) Schematic protocol of malaria egress media purification. Schizonts are purified from a synchronous asexual blood stage culture and exposed to compound 2, a schizont rupture inhibitor. After incubation, late schizonts are resuspended in endothelial media for simultaneous rupture in defined concentrations. Soluble factors are purified by centrifugation, and the resulting supernatant is used for downstream endothelial assays. (**B and C**) GO term over-representation analysis of (**B**) upregulated and (**C**) downregulated GO biological processes. Shown are the 75 most significant processes (adjusted p-value < 0.05, log_2_FC > ± 0.1). Node connections indicate shared genes between GO terms, and node clusters are manually annotated. (**D, E and F**) Heatmaps display log_2_ fold changes after 8- and 24-hour egress product exposure compared to media controls of key genes in (**D**) steroid biosynthesis pathways, (**E**) cholesterol metabolism and (**F**) spliceosome pathways. Asterisks indicate significant DEseq2 expression (adjusted p-value < 0.05) for one (*) or both (**) time points. (**G**) Distribution of alternatively spliced events by type. Plotted event numbers are differentially spliced (ΔPSI ≥ 15) between the ETS-iBMEC basal condition and *P. falciparum-*infected RBC at 24h. Alternative 3’/5’ ss, alternative splice-site acceptor/donor selection. (**H**) Predicted impact of alternatively spliced events on the proteome based on data from VastDB. Many of the events are predicted to disrupt ORFs (e.g., frameshift and/or premature termination codon) or to generate alternative protein isoforms.

Shared dysregulated GO term processes included upregulation of sterol and cholesterol synthesis pathways (Fig. 5B, D, E, Fig. S5C). Other upregulated processes in ETS-iBMEC included circulatory regulation and synaptic organisation, with the latter including endothelial transporters important for barrier regulation (AMPAR, NMDAR, KAR, CACN) and angiogenesis-related transcripts (*NTN1, SEMA3, SEMA4, SEMA7, L1CAM*) (Table S2). A common downregulated process among the three cell types, with special relevance in ETS-iBMEC, was RNA splicing (Fig. 5C), along with other terms associated with RNA processes and translational regulation (Fig. 5C, Fig. S5C). Spliceosome transcripts showed early decreased expression after 8 hours, sustained throughout the 24-hour experiment (Fig. 5F). Because alterations in pre-mRNA processing can result in profound changes in the endothelial cell biology, we then focused our analysis on alternative splicing after exposure to infected RBC products. Using vast-tools (*36*), we identified 692 events (about 7%) that were differentially spliced between the two conditions (more than 15% of a particular mRNA alternatively spliced; ΔPSI ≥ 15) (Table S3). The most common events included alternative exon splicing (37.7%) and intron retention (28.3%) (Fig. 5G). Based on data from VastDB, many of the detected events are predicted to disrupt the open reading frame (51.3%) or create alternative protein isoforms (30.7%) (Fig. 5H). Moreover, 112 events had substantial changes in splicing (ΔPSI ≥ 25). Among them were proteins responsible for autophagy (ATG16L2, RB1CC, SQSTM1), with important functions in cytoskeleton dynamics (Tropomyosin 1), and a histone deacetylase regulating inflammatory gene expression (HDAC5/7/8) (Table S3). These data suggest that infected RBC products cause profound changes in ETS-iBMEC homeostasis, including alterations in transcription and/or potential changes in alternative splicing.

### *P. falciparum-*egress products disrupt the ETS-iBMEC barrier and alter the transcription of endothelial junctional, cytoskeletal, and angiogenic pathways

Endothelial barrier disruption plays an important role in severe malaria, especially in complications such as cerebral malaria, characterised by severe brain swelling and blood-brain barrier disruption (*37–39*). Multiple downregulated processes in the transcriptomic analysis were suggestive of changes in barrier integrity, including cell migration, actin organisation, substrate adhesion, and endothelial development (Fig. 5C). To determine whether these changes would lead to endothelial barrier disruption in ETS-iBMECs, we assessed trans-endothelial resistance through real-time impedance-based measurements (xCELLigence). ETS-iBMECs were exposed to 1.25, 2.5, and 5×10⁷ infected RBCs/mL—concentrations corresponding to the simultaneous egress of *P. falciparum*-infected RBCs at parasitemias of 0.25%, 0.5%, and 1%, respectively (*37*). All conditions tested led to a decrease in the cell index, indicative of compromised endothelial integrity. Despite high variability, a plateau in endothelial breakdown was observed at 2.5×10⁷ ruptured infected RBCs/mL (Fig. 6A, B). Overall, our results reveal that ETS- iBMECs respond to the addition of *P. falciparum-*egress products through endothelial barrier disruption.

**Fig. 6.**
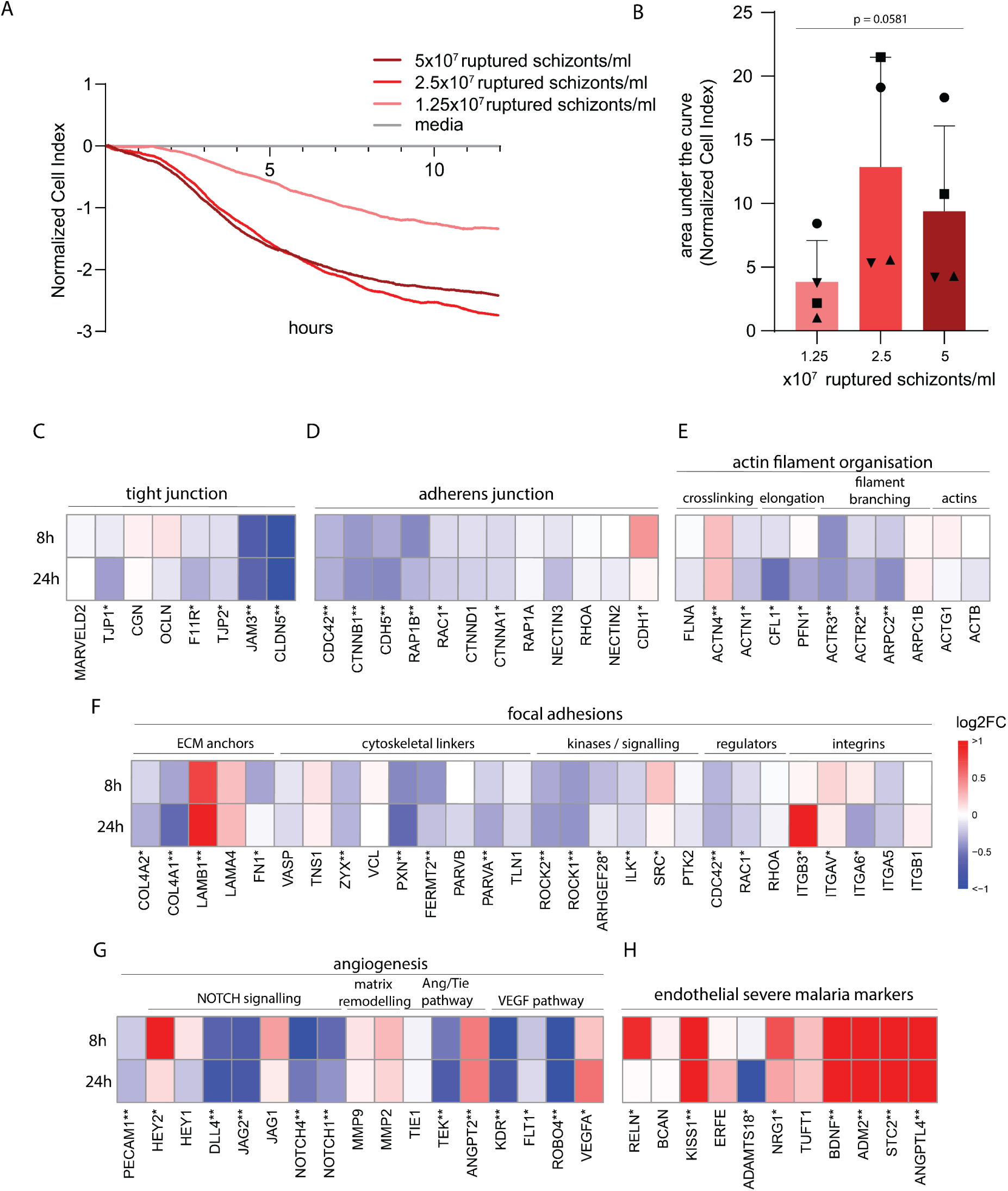
Transcriptional changes induced by *P. falciparum* egress products in ETS-iBMECs across barrier-associated gene categories. (**A**) Representative record of normalized cell index from real-time xCELLigence trans-endothelial impedance measurements of ETS-iBMECs exposed to egress media in different concentrations. **(B)** Resulting area under the curve analysis for 12 hours for four independent cell preparations (one-way ANOVA). (**C-H**) Heatmaps display log_2_ fold changes after 8- and 24-hour egress product exposure, compared to media controls, for key genes involved in the architecture of tight junctions (**C**), adherens junctions (**D**), actin filaments (**E**), focal adhesions (**F**), a selection of key angiogenesis genes (**G**), and endothelial severe malaria marker genes (**H**) (*40*). Asterisks indicate significant DESeq2 differential expression (adjusted p-value < 0.05) for one (*) or both (**) time points.

A targeted transcriptomic analysis of endothelial junctional genes revealed a coordinated and pronounced downregulation of adherens and tight junction markers, highlighting a potential mechanistic axis of endothelial barrier destabilisation in response to malaria products in ETS-iBMEC (Fig. 6C-D). Notably, key tight junction members, such as *CLDN5* (Claudin-5), *F11R* (JAM-1), and *JAM3*, along with tight junction adaptors *TJP1* (ZO-1) and TJP2 (ZO-2), were significantly downregulated, some early upon exposure and others at later timepoints. Decreased expression of adherens junctional transcripts, *CDH5* (VE-cadherin), *CTNNB1* (β-catenin), and *CTNNA1* (α-catenin), as well as other junctional components such as *PECAM1* (CD31, Fig. 6G), was also found. In addition to the decrease in cell-junction transcripts, *P. falciparum* egress products downregulate key genes involved in actin dynamics. While actin isoforms (*ACTB, ACTG1*) remained stable, components of the Arp2/3 complex (*ARPC2*, *ACTR2*, and *ACTR3*), involved in filament branching, were downregulated early upon exposure (Fig. 6E). Furthermore, we observed early (8h) and sustained (24h) transcriptional repression of most focal adhesion-associated genes including *ZYK* (zyxin), *PXN* (paxillin), and *TLN* (talin), whereas others such as *VCL* (vinculin) remained unchanged. Key regulators of focal adhesion formation and stabilisation (ROCK1, ROCK2), along with late downregulation of integrins, were also revealed. This occurred simultaneously to disruption in extracellular matrix secreted proteins, including *COL4A1* and *COL4A2* downregulation and *LAMB1* upregulation (Fig. 6F). We also identified alterations in key vascular homeostatic and angiogenic processes, including members of the VEGF pathway (*VEGFA*, *ROBO4*, *FLT1*, and *KDR),* Notch signalling (*NOTCH1, NOTCH4*, *JAG2*, and *DLL4)*, and Angiopoietin/Tie axis (*ANGPT2* and *TEK*) (Fig. 6G).

Notably, the downregulation of junctional, cytoskeletal and focal adhesion transcripts was more pronounced and robust in ETS-iBMEC than in primary brain ECs (Fig. S6). However, ETS-iBMEC did not show overexpression of viral processes and interferon response, including the JAK-STAT pathway, a key pathogenic response predominantly found in primary brain ECs in a 3D-blood brain barrier (*26*) (Fig. S5C). Nevertheless, we identified additional important transcriptional dysregulations in primary endothelial cells that have been recognised as biomarkers of clinical severe malaria (28). Of the selected markers, seven showed significant upregulation at one or both time points, with the strongest induction observed for ANGPTL4, STC2, ADM2, BDNF, and KISS1 (Fig. 6H). Overall, these results reveal that *P. falciparum*-infected RBC products profoundly affect vascular homeostatic pathways and highlight the use of ETS-iBMECs as a relevant model for malaria pathogenesis.

## Discussion

Research on severe malaria has been hampered by the lack of an animal model that recapitulates the accumulation of *P. falciparum-*infected RBC in the brain microvasculature. As an alternative, most *in vitro* pathogenesis models have used immortalised or primary cells, which present inherent limitations such as loss of cell identity and batch-to-batch variability (*26, 27, 34, 41–43*). The use of iPSC-differentiated endothelial models for malaria research emerges as a promising alternative, allowing the future development of patient-specific isogenic models. Although iPSC-based models have recently been applied to model cerebral malaria (*12*), challenges to faithfully recapitulate endothelial identity remain the main limitation in the stem cell bioengineering field (*8*). Here, we have generated a stable iPSC transgenic line that directs endothelial differentiation of iBMECs through DOX-induction of three key ETS transcription factors: ETV2, ERG, and FLI1. We confirmed ETS-iBMECs as a valid model to study malaria pathogenesis through binding studies by perfusing *P. falciparum*-infected RBCs under flow, as well as transcriptomics. The newly developed models reveal that ETS-iBMECs form elevated trans-endothelial barriers that are susceptible to breakdown upon exposure to parasite products, and are associated with downregulation of tight and adherens junctional markers, focal adhesion, and cytoskeletal markers, along with numerous differential splicing events.

The essential role of ETS transcription factors in endothelial development is well established (*8, 13, 15, 17, 44–47*), among which ETV2 particularly drives the emergence of vasculogenic, generic endothelial cells (*48, 49*). Overexpression of ETV2 has been widely used to promote endothelial differentiation, including electroporation (*13*), lentiviral transduction (*8, 15–17*) or more recently with the generation of stable lines through piggyBac-mediated integration (*45–48, 50, 51*). Our study, to our knowledge, is the first to use a stable, transgenic iPSC line enabling inducible upregulation of three ETS-factors, thereby robustly promoting endothelial identity. Consistent with previous studies (*8, 13, 15, 17, 44–48*), DOX induction in ETS-iBMECs strongly directed the differentiation toward the endothelial lineage, as shown by increased expression of endothelial markers and ultrastructural morphology, while maintaining enhanced brain-*like* barrier function. Our approach not only streamlines experimental procedures but also allows for the selection of a cell line with three stably integrated inducible transcription factors, including ETV2, to further promote endothelial differentiation.

Although ETS-induction is not intended to generate brain-specific endothelial cells, ETS- iBMECs markedly outperformed primary brain ECs in terms of barrier function in 3D microvessels. The decline in barrier function commonly observed in primary brain EC *in vitro* is often attributed to the loss of tight junctional features [12,27-29], supported by the reduced expression of ZO-1 (*TJP1*) and occludin (*OCLN*) transcript levels found in primary cells in our transcriptomic analysis. Conversely, ETS-iBMEC displayed increased expression of these markers and showed minimum permeability to both low and medium-molecular-weight dextran tracers in a 3D microvascular BBB model, while keeping endothelial ultrastructural morphology, including thin apical-basal height, loss of desmosomes, and lack of microvilli. Future iterations of the model to improve the differentiation toward brain-specific lineage, relevant for cerebral malaria studies, could incorporate FOXF2 and ZIC3, as the recent inducible expression of these brain-specific transcription factors via lentiviral transduction has shown enhanced brain identity and strengthening of barrier properties (*52*). Alternatively, the addition of Wnt activators and TGFβ inhibitors has also recently been shown to further promote brain endothelial specification (*45*).

ETS-iBMECs were incorporated in a 3D BBB microvascular model, increasing the physiological complexity through co-culture with astrocytes and pericytes. While this is a proof of concept, in order to implement ETS-iBMECs in more physiologically complex systems, we employed higher throughput 2D microfluidic and real-time impedance-based barrier assays to model *P. falciparum* pathogenesis. These included *in vitro* studies characterising *P. falciparum-*infected RBC sequestration, one of the main hallmarks of severe and cerebral malaria (*53–57*). Perfusion of *P. falciparum-*infected RBC through commercial microfluidic flow chambers revealed strong binding affinity to ETS-iBMECs, supporting their relevance as an alternative model of parasite cytoadherence. The parasite lines used in this study expressed PfEMP1 variants that are highly enriched in severe and cerebral malaria patients (*58–60*). The dual CD36-ICAM 1 binder, ITGICAM1, showed increased binding to ETS-iBMECs compared to iBMEC-*like* and primary brain ECs. ETS-iBMECs expressed the highest levels of ICAM1 under resting conditions and low levels of CD36. Therefore, it is likely that the increased binding of ITGICAM1 is mediated by ICAM1, given its strong affinity to the receptor (*61*). The EPCR binder, IT4var19, also showed increased binding to ETS-iBMECs compared to the other two cell types. Although this phenotype cannot be explained through our transcriptomic analysis, as the expression of *PROCR* (EPCR) was higher in primary brain EC, post-translational modifications or differences in receptor trafficking to the surface may account for the non-significant increase in binding in ETS-iBMECs. The pronounced binding of *P. falciparum* to ETS-iBMECs highlights this model’s relevance for studying host–parasite interactions in severe malaria. Future studies could include additional parasite lines with a dual EPCR–ICAM1 binding phenotype (*61, 62*) or samples from patients with severe and uncomplicated malaria to gain deeper insights into disease mechanisms, as cerebral malaria isolates have shown increased affinity for primary brain endothelial cells (*53*). Furthermore, future strategies to gain deeper molecular insights into host–parasite interactions could include knocking out EPCR and ICAM1 in the parental line or introducing point mutations. Therefore, ETS-iBMEC represent an experimentally tractable and physiologically relevant platform to test intervention strategies aimed at reducing *P. falciparum* cytoadhesion (*61, 63*),

In fatal cases of cerebral malaria, the blood-brain barrier is disrupted, as showing by magnetic resonance imaging scans revealing severe brain swelling (*37, 38*), and by signs of fibrinogen leakage seen in post-mortem samples (*64*). Furthermore, brain vasogenic oedema is not exclusive to severe disease and has also been observed in cases of uncomplicated malaria (*39*). Our previous work, along with others, has shown that *P. falciparum* egress in close proximity to endothelial cells is a major driver of barrier breakdown in primary brain EC (*26, 27, 42*). Notably, this phenotype was recapitulated in ETS-iBMECs using impedance-based assays. The disruption was accompanied by a global and homogenous downregulation of almost all transcripts belonging to tight and adherens junctions, among them occludin and ZO-1, previously found decreased in histopathological labelling of cerebral malaria from paediatric (*65*) and adult samples (*66*). Downregulation of vinculin, a focal adhesion junctional protein, was also found in patient post-mortem samples. Although we did not observe a significant downregulation of vinculin upon ETS-iBMEC exposure to parasite products, a significant decrease in expression of almost all other structural and signalling focal adhesion proteins was found. Remarkably, such robust and global downregulation of key regulatory barrier pathways by malaria parasites or products has not been found in previous primary 2D (*42*) or in a 3D endothelial-only model (*43*). Alterations in some key genes in these pathways have been identified in our recent *in vitro* primary 3D-BBB model co-cultured with astrocytes and pericytes (*26*), but at a much lower global extent than ETS-iBMECs. This may be a consequence of the greater maturation of endothelial junctions in ETS-iBMECs compared to primary ECs. Alternatively, the global and coordinated response in ETS-iBMECs could result from a more homogenous cellular identity, compared to the heterogeneity observed in primary brain ECs, which arises from their diverse anatomical origin across brain regions and vascular segments (*67*), such as arterioles, capillaries, and venules.

Despite these differences in the response to *P. falciparum* between primary brain ECs and ETS-iBMECs, we also identified similarities, such as important transcripts for malaria pathogenesis that were later classified as serum biomarkers of disease (*40, 42*). Furthermore, the ETS-iBMEC model reproduced alterations in angiogenic pathways important for disease, such as VEGF (*68*) and the angiopoietin/tie2 axis. A disruption of the angiopoietin/Tie2 axis has also long been recognised as a biomarker of cerebral malaria (*69–72*), and targeting this pathway has been proposed as a potential therapeutic (*27, 73*). Furthermore, higher concentrations in plasma of VEGF have been found in uncomplicated (*74*) and in the brain of cerebral malaria patients at sites of *P. falciparum-*infected RBC sequestration (*75*) and are strongly associated with patient fatality (*68*). Overall, the ability of ETS-iBMEC to replicate all these relevant pathways of severe disease underscores their ability of iPSC-derived cells to reproduce established relevant molecular signatures of disease.

Nonetheless, limitations remain. The differentiation protocols require specialised training and are laborious, and although ETS-iBMECs can be incorporated in complex 3D microvessel models, the low throughput hampers their application to infection research in complex *in vitro* systems. Furthermore, we found that ETS-iBMECs do not activate processes associated with the response to viruses upon challenge with *P. falciparum-* infected RBC products. We have recently shown the importance of the IFN type I and JAK-STAT pathway in *P. falciparum-*mediated barrier disruption in a primary 3D-BBB model. This model showed that ruxolitinib, a JAK-STAT inhibitor, could prevent the loss of vascular integrity, a finding consistent with human clinical trials that revealed the efficacy of ruxolitinib at preventing vascular damage during controlled malaria infections, as well as other parameters associated with severity (*42, 76–78*). Therefore, future improvements in endothelial differentiation approaches should aim to streamline protocols to develop complex 3D assays and enhance innate immune responses in iPSC-derived cells.

Nevertheless, the ETS-iBMEC model recapitulates key pathogenic events of severe malaria and opens new avenues for studying its pathogenesis. Future applications of stem-cell technologies could include the generation of models with patient-derived iPSCs, the generation of isogenic multicellular immunocompetent models that could include other perivascular cells, as well as peripheral or tissue resident immune cells from a single donor, given the recently identified importance of T-cells in cerebral malaria (*79, 80*). Other future benefits could include the generation of CRISPR-Cas screenings that could shed light on the molecular mechanisms that drive severe malaria disease pathogenesis (*81*). Overall, ETS-iBMEC-based models establish the value of stem cell-derived systems for *in vitro* studies of severe malaria, paving the way for future advances in disease modelling, therapeutic testing, and mechanistic understanding.

## Methods

### iPSC culture and differentiation

Human iPSCs (IMR90-4, WiCell Research Institute, Inc.) were maintained on Matrigel-coated dishes with mTeSR1 (Stemcell Technologies) maintenance medium. The DOX-inducible transgenic cell lines for the upregulation of human ETV2, ERG, and FLI1 were generated as follows. Each gene sequence was cloned into the pDONR P5-P2 entry vector, while a modified Tet-One system (Takara Bio) was cloned into the pDONR P1-P5r vector. These entry vectors were then recombined into a piggyBac destination vector containing an antibiotic resistance cassette using the MultiSite Gateway Pro system (ThermoFisher) (*82*). The resulting constructs were electroporated into IMR90-4 iPSCs using the Amaxa 4D-Nucleofector protocol (Lonza), followed by antibiotic selection of the transgenic cells. To assess silencing effects on Tet-One upregulation during differentiation, the same design has been generated with fluorescent reporter proteins. Differentiation has been performed as described (*7*). In brief, iPSCs were seeded as single cells on Matrigel-coated 6-well plates in mTeSR1 with Y-27632 and maintained for two further days in mTeSR1 only. Equally distributed colonies were exposed to unconditioned medium (UM) for six days, containing: 39.25 mL DMEM/F12 (Invitrogen) with 10 mL Knockout Serum Replacement (Gibco), 0.25 mL Glutamax (Invitrogen), 0.5 mL non-essential amino acids (Invitrogen) and 0.35 μL ß-mercaptoethanol (Sigma) at 5% O_2_. Afterwards, endothelial expansion and transcription factor upregulation were performed for two days in human endothelial serum-free medium (hESFM, Invitrogen), with 1% plasma-derived serum (Sigma), 20 ng/mL hFGF (R&D Systems), and 10 μM retinoic acid (Sigma) for iBMEC-*like* cells. ETS-iBMEC differentiation was promoted through the addition of 100 ng/mL doxycycline. The generated cells were accutase-released, passaged for one hour on collagen-fibronectin-coated wells, washed and used for further assays. TNFα stimulation occurred at 10ng/mL for 18 hours. Mycoplasma tests were performed biweekly.

### Primary cell culture

Primary human brain microvascular endothelial cells (HBMEC, ACBRI 376, Lonza) were cultured in microvascular endothelial cell growth media supplemented with 5% FBS and endothelial supplements (EGM2-MV, Lonza). Primary human astrocytes (HA, ScienCell) were cultured in basal medium supplemented with 2% FBS, 1% Pen-Strep and 1% astrocyte growth factor. Primary human brain vascular pericytes (HBVP, ScienCell) were cultured in basal medium with 2% FBS, 1% Pen-Strep and 1% pericyte growth factor. For culture, all primary cell types were grown in monolayers on Poly-L-lysine-coated flasks with media changes every two days up to 90% confluency. Cells were passaged with Trypsin/EDTA up to passage number 8. Mycoplasma tests were performed biweekly.

### P. falciparum culture

*P. falciparum* parasite lines HB3var03, IT4var19 (EPCR binding) (*83*) and ITGICAM1 (ICAM1 binding) (*84*) were cultured in human O+ erythrocytes in complete media (Gibco RPMI 1640 medium, 10% human type AB-positive plasma, 0.4 mM hypoxanthine, 5 mM glucose, 26.8mM sodium bicarbonate, 1.5 g/L gentamicin), in culture flasks gassed with 90% N_2_, 5% CO_2_ and 1% O_2_. Parasites were synchronised twice a week with 5% sorbitol (Sigma) for the synchronisation to ring stages, and 40 mg/mL gelaspan (Braun) for the purification of trophozoites.

### 3D-BBB fabrication

The generation of 3D microvessels was performed as previously described for blood-brain barrier models (*26, 85*) with either of the three different endothelial cell sources: primary HBMEC, iBMEC-*like* cells and ETS-iBMEC, in combination with primary HA and HBVP. Briefly, type I collagen was isolated from rat tails and dissolved in 0.1% acetic acid (Sigma) at 15mg/mL and then diluted to 7.5 mg/mL in endothelial media supplemented with 1% astrocyte and pericyte growth factor (AGF and PGF). Primary 7.5x10^5^ HA/mL and 3.2x10^5^ HBVP/mL were added to the neutralised solution before fabrication. The microvascular model was manufactured in a process combining soft lithography and injection moulding as previously described (*85*). The collagen solution containing HA and HBVP was injected on the top and bottom polymeric PMMA jigs previously treated with O_2_ plasma. The top part was obtained by negative impression of a polydimethylsiloxane (PDMS) 2-channel pre-patterned microfluidic network. The bottom part was generated through patterning with a flat stamp. Both parts were left for gelation for 1 hour at 37°C, followed by removal of the PDMS stamps, and top and bottom jigs assembly. For microvessel formation, 7x10^6^ HBMEC/mL were resuspended in EGM2-MV (Lonza) with 1% AGF and PGF and seeded in the device by gravity-driven flow. iBMEC-*Iike* and ETS-iBMEC were seeded at 23x10^6^ cells/mL in hESFM with 1% PDS, AGF and PGF after previous coating with collagen/fibronectin (400/100 ng/mL) for 45 minutes at 37°C. The microvessels were fed with the respective media twice a day. ETS-iBMEC had continuous DOX exposure until further experimental use.

### Immunofluorescence assays

Monolayers and 3D microvessels were fixed in 4% paraformaldehyde for 20 minutes and washed with PBS. The cells were exposed to Background Buster (innovex Bioscience) for 30 minutes and blocked/permeabilised with 2% BSA and 0.1% Triton X-100 in PBS. Primary antibodies for CD31 (BD Pharming, 560983, 1:100), EPCAM (Biolegend, 324202, 1:100), ZO-1 (Invitrogen, 40-2200, 1:100), β-catenin (Sanat Cruz Biotech, 59737, 1:100), PAR-1 (R&D System, FAB3855G, 1:100), ICAM1 (Abcam, ab20, 1:100) and Phalloidin (Invitrogen A22287, 1:200) were added in blocking solution overnight. Secondary antibodies and DAPI (ThermoFisher, 8 μg/mL) were added after washing for 1 hour. After additional washes in PBS, monolayers and microvessels were kept and imaged in PBS. Imaging was performed on an LSM980 Airyscan 2 (Zeiss) and processed using the ZEN imaging software (Zeiss) and Fiji (ImageJ).

### Dextran permeability assays

The barrier function of 3D microvessels was tested by permeability assays with low molecular tracers after 4 days in culture in a 37°C microscope environmental chamber on an LSM980 Airyscan 2. Perfusion of 40 kDa (10μM, FITC labelled) and 10 kDa (2 μM, AF647 labelled) fluorescent dextran was performed at a flow rate of 6.5 μL/min, reflecting a wall shear stess of 1 dyn/cm^2^ with a withdrawing syringe pimp (Harvard Apparatus, PHD 2000). The z-focus was adjusted to the central plane of the vessel, and acquisition intervals of 30 seconds were captured. For quantification, timepoint t_0_ was set after full filling of the microvessels with the fluorescent tracer and t_1_ after 2.5 minutes of perfusion. Equally sized ROIs (150 μm x 150 μm) were set along the microvessel wall with an equal distribution on the vessel (150 μm x 30 μm) and hydrogel area (150 μm x 120 μm). Apparent microvascular permeability in cm/s was calculated by integrating fluorescence intensity measurements of the set regions between the two timepoints into the following equation: P = (1 / Δt) (V_gel_ / A_v_) ((I_gel1_ - I_gel0_) / (I_v0_ - I_gel0_)), in which Δt is the time interval between t_0_ and t_1_, V_gel_ is the volume of the collagen matrix, A_v_ the lateral vessel surface, I_gel1_ - I_gel0_ is the difference in measured intensities in the gel area between t_1_ and t_0_, and I_v0_ is the fluorescent intensity in the vessel at t_0_.

### Barrier integrity assays by xCELLigence

The ETS-iBMECs were grown on collagen-fibronectin-coated 96-well PET E-plates (Agilent) during real-time cell-index measurements in the xCELLigence RTCA SP reader. Cell attachment and barrier formation was assessed for three days until the cell-index plateaued with a media change every two days. To test for dose-dependent effects on barrier integrity of the egress media, concentrations equivalent to 5, 2.5 and 1.25x10^7^ ruptured infected RBCs were adjusted with the stock egress media in hESFM with 1% PDS and DOX. After application, the cell index was measured in 15-second intervals in parallel to a media control. All conditions were applied in triplicate, and each experiment was performed with independent ETS-iBMEC differentiations and individual parasite egress media purifications. For analysis, the cell index progression was set to 0 after sample application and sedimentation. The averaged triplicates of the cell index were normalised to the cell index of the media-only control. Area under the curve analysis was performed using GraphPad Prism (version 10.0.2) for experimental times of 12 hours.

### Transmission electron microscopy

The 3D microvessels were fixed with 2% PFA and 2.5% GA in endothelial media by perfusion for 30 minutes, followed by two media washes. The device was disassembled, the hydrogel removed from the PMMA jig and cut into small areas. These areas were processed with a secondary fixative (2% PFA, 2.5% GA, 0.25 mM CaCl_2_, 0.5mM MgCl_2_, 5% sucrose in 0.1% sodium cacodylate in water) over night at 4°C and post-fixed in a reduced osmium solution (1% OsO_4_, 1,5% K3F3(III)(CN)6 in 0.065M sodium cacodylate buffer) six times. Post fixation, the hydrogels were incubated for two hours in OsO_4_ at 4°C and washed with water six times. The samples were dehydrated in steps of 30, 50, 80, and thrice with 100% ethanol in a PELCO Biowave Pro microwave processor (Ted Pella, Inc.) containing a SteadyTemp Pro and a ColdSpot set to 4°C, with 40 seconds at 250 W each. Infiltration was performed in serial steps of 25, 50, 75, 90, and twice with 100% EPON 812 hard epoxy resin in acetone, assisted by the microwave (3 minutes each at 150 W under vacuum) and a final infiltration step in 100% EPON 812 hard epoxy resin overnight at room temperature. In the embedding mould, samples were positioned with the microvessel luminal axis at a 90° angle to the cutting surface to cut transversal sections of the vessel, and then polymerised at 60°C for 48 hours. Microvessel pieces for imaging were randomly selected, and thin sections (70 nm) were retrieved on an ultramicrotome (UC7, Leica Microsystems), collected on formvar-coated slot grids and post-stained in uranyl acetate and lead citrate. Tile montages (12,000x) were acquired on a JEOL JEM 2100 plus at 80 or 120 keV using SerialEM. Montages were processed using IMOD’s Blend Montages function and Fiji (ImageJ).

### *P. falciparum var* typing and perfusion

For parasite binding assays, *P. falciparum* lines It4var19 and ITGICAM-1 were previously panned and monitored for the respective PfEMP1 variant expression. Briefly, parasites are synchronized with 5% sorbitol and the RNA of ring stage parasites (0-22 h) post-invasion is extracted with Tryzol LS and an RNAeasy kit, and reverse transcribed with TaqManTM Reverse Transcription Reagents (Applied Biosystems, N8080234). The expression of the *var* repertoire is monitored with an IT4 primer repertoire (*86*) with LightCycler 480 SYBR Green I Master (Roche, 04707516001) in a LightCycler 480 II (Roche) following published amplification conditions. Relative transcription of *var* genes was normalized to the housekeeping control seryl-tRNA-synthetase (STS; PF07_0073) and the expression was represented as relative gene expression = 2(- ΔCT). For *P. falciparum* binding assays, the synchronised parasite strains were grown to high parasitemia (5-10%) and purified using Percoll (Cytiva) or LD columns (Miltenyi Biotec) to obtain cultures >60% parasitemia. Infected RBCs were stained with PKH67 fluorescent membrane dye (Sigma) and gravity perfused three times at 1x10^7^ infected RBC/mL through endothelial μ-slides VI 0.1 (ibidi) channels previously seeded three days before with iBMEC-*like*, ETS-iBMEC and primary brain EC. After parasite perfusion, unbound parasites were washed out with media for 10 minutes, and the endothelial channels were fixed with 4% PFA in PBS for 15 minutes and washed with PBS. After fixation, endothelial nuclei were stained with DAPI (8 μg/mL), and the channels were imaged using confocal microscopy on an LSM980 Airyscan 2 (Zeiss) and processed using Fiji (ImageJ). Up to nine ROIs of equal size were selected per channel and bound parasites quantified using Fiji.

### Parasite egress media preparation

Parasite egress products were purified by tightly synchronising *P. falciparum-*infected RBC HB3var03 parasites into a 6-hour window. Late-stage parasites, approximately at 44 hpi, were enriched to 5x10^7^ infected RBC/mL using Percoll purification, and incubated in complete media with 1 µM compound 2, which inhibits schizont egress (a reversible PKG inhibitor kindly provided by Michael Blackman, The Francis Crick Institute). After 5 hours, simultaneous schizont rupture was induced by resuspending parasites in endothelial media without compound 2 at a concentration of 10^8^ schizonts/mL overnight at 30 rpm on a shaker. The total rupture efficiency was assessed by a hemocytometer count, and Giemsa blood smear and supernatants were recovered after spinning at 1000 rpm and snap frozen in liquid nitrogen. For experimental use, the concentration of rupture released factors was adjusted to the equivalent of 5, 2.5 and 1.25x10^7^ ruptured schizonts/mL, which represents the physiological simultaneous release during approx. 1% parasitemia within one asexual blood cycle (*37*).

### Bulk RNA-sequencing and analysis

For transcriptional analysis, bulk RNA sequencing data were generated from primary brain EC, iBMEC-*like* cells, and ETS-iBMEC. Cells were cultured as 2D monolayers for three days on 24-well plates and exposed to egress products from ruptured schizonts (5x10^7^ schizonts/mL) or a media control. The experimental design included exposure durations of 8 and 24 hours to examine transcriptional responses over time. Samples were fixed with RLT buffer, and RNA was extracted using the RNeasy Mini kit (QIAgen), including DNAse treatment. Quality and RNA concentration were assessed by TapeStation (Agilent). Individually barcoded stranded mRNA-seq libraries were prepared from high-quality total RNA samples (∼100 ng/sample) using the New England Biolabs NEBNext RNA Ultra II Kit combined with the polyA-enrichment module in 12 PCR cycles on a liquid handling robot (Beckman i7). The resulting libraries were pooled in equimolar amounts and loaded on the Illumina sequencer NextSeq 2000 platform, sequencing unidirectionally, generating 1980 million reads, each 80 bases long.

Raw reads were assessed for quality using FastQC (version 0.12.1). Transcript quantification was performed using Salmon (v1.10.3) with an index built from the Ensembl GRCh38 reference transcriptome (*87*). Transcript-level abundances from Salmon were imported into R and summarised to gene level using the tximport package (v1.32.0) (*88*), with transcript-to-gene mappings derived from the corresponding Ensembl GTF annotation. Only protein-coding genes were retained for further analysis, as determined by annotation with org.Hs.eg.db (v3.19.1). Genes with low read counts were filtered out (<10 counts in at least one sample) before differential expression analysis, which was performed using DESeq2 (v1.44.0) (*89*). Log fold change shrinkage was applied with apeglm (v1.26.1) for more stable effect size estimation (*90*). P-values were adjusted for multiple testing using the Benjamini-Hochberg procedure. Genes with adjusted p-value < 0.05 and |log₂ fold change| > 0.1 were considered significantly differentially expressed. Log_2_ fold changes (LFCs) for selected gene sets of interest were extracted from DESeq2 differential expression results. Heatmaps were generated using the pheatmap package (v1.0.12) (*91*). GO over-representation analysis of differentially expressed genes was performed using the clusterProfiler package (v4.12.6) (*92*). Ensembl gene identifiers were mapped to Entrez IDs using org.Hs.eg.db. GO over-representation was carried out for biological process terms. GO term maps (emapplots) were generated with enrichplot (v1.24.4) to display similarity networks of enriched processes, showing clusters of functionally related terms among up- or downregulated gene sets with a semantic similarity score ≥ 0.15 (*93*). Volcano plots were visualised using the EnhancedVolcano package (v1.22.0) (*94*). Endothelial and epithelial gene sets were highlighted using curated gene lists from endothelial (red) or epithelial (blue) GO terms. Marker genes from each group with a described endothelial or epithelial function were labelled. Variance stabilising transformation was performed to generate normalised gene expression heatmap visualisations. A curated set of genes of interest was selected, and their expression values were averaged for each condition. The centered, normalised expression values were visualized as heatmaps using the pheatmap package, with clustering performed on columns (conditions).

### Detection of alternative splicing events and analysis

Alternative splicing profiles have been generated using vast-tools, which aligns and processes raw RNA-seq reads to derive PSI (percent spliced in) values for all types of AS events. *Homo sapiens* (hg38, Hs2), with Ensembl v88 scaffold annotation was used for event detection. Differentially spliced events between the media and the 24 hour timepoint condition were considered when ΔPSI was at least 15. The potential impact of splicing on the proteome was based on data from VastDB (*36*).

### Statistical analysis

All statistical tests were performed with GraphPad Prism. Shapiro-Wilk test has been used to test normality. One-way ANOVA was used for testing differences between groups in normally distributed data. For nonparametric data, Mann-Whitney U test has been performed to test differences between two groups, and Dunn’s multiple comparison test was used to compare multiple pairs within the dataset.

## Data availability

RNA-seq datasets generated and used in this study are available in the ArrayExpress database under the accession code E-MTAB-14937.

## Author contributions

FK and MB conceived the work. FK, HF, RL, LP, ME and MB designed the experiments. FK, HF, LP, BG, and DC performed the experiments. FK analysed the experimental data under the guidance of ME and MB. PP analysed splicing events under the supervision of VT. AB gave guidance on RNA-seq analysis performed by FK. FK and MB wrote the original draft of the manuscript. All authors contributed to the manuscript revision, editing and suggestions. FK, ME and MB contributed to project supervision, administration and funding acquisition.

## Supporting information

Table S1 GO enrichment iBMEC-like vs ETS-iBMEC

Table S2 GO enrichment ETS-iBMEC egress media

Table S3 Differential alternative splicing ETS-iBMEC

## Acknowledgments

We want to thank all members of the Bernabeu and Ebisuya labs for their supportive discussion and critical feedback. We want to thank Mitsuhiro Matsuda for the support and mentoring with the stem cell technologies. Furthermore, we want to thank Jorge Trojanowski for input on the RNA-seq experimental preparation and analysis, Jaroslaw Sochacki, Jorge Lázaro, and Laura Batlle for feedback on iPSC culture, Michael Blackman (The Francis Crick Institute) for kindly providing compound 2 and Knut Woltjen for providing the piggyBac vector.

We thank Vladimir Benes and the Genomics Core facility (Gene Core) at EMBL, the Genomics Core facility at the Universitat Pompeu Fabra (UPF), Charles Girardot and the Genome Biological Support (GBCS) at EMBL for their support in RNA sequencing and the Electron Microscopy Core Facility (EMCF) at EMBL for their support with EM imaging. This work was supported by the European Research Council (ERC), the European Union’s Horizon 2020 research and innovation program (Grant agreement No. 948088 to MB and No. 101002564 to ME), the Deutsche Forschungsgemeinschaft (DFG, German Research Foundation) under Germany’s Excellence Strategy - EXC 2068 - 390729961 - Cluster of Excellence Physics of Life of TU Dresden, the Alexander von Humboldt Foundation, and the Marie Skłodowska-Curie Actions COFUND (847543) with additional funds from the EMBL core program and the EMBL Infection Biology Transversal Theme.

---

**Fig. S1.**
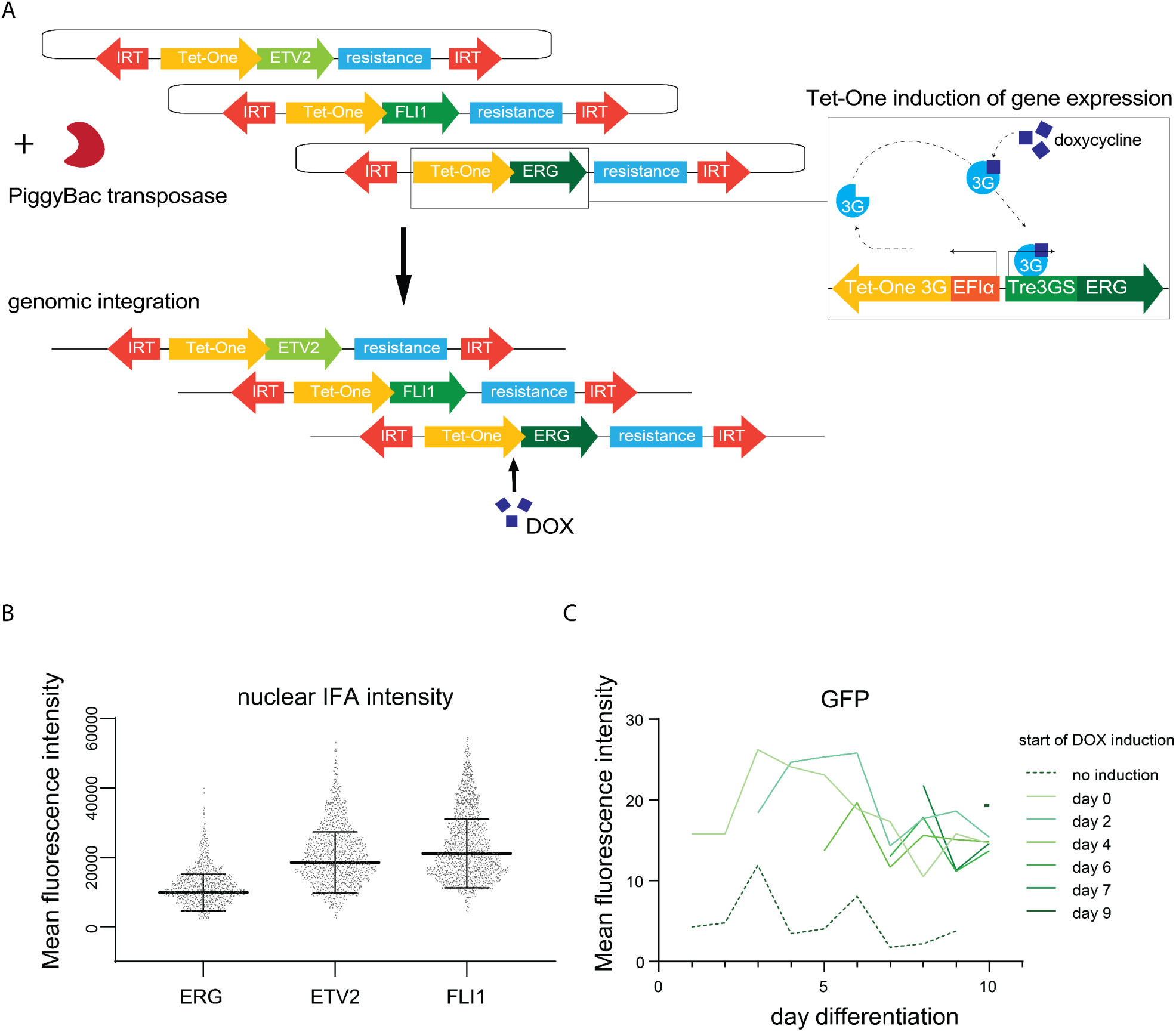
ETS-transcription factor upregulation in ETS-iBMEC. (**A**) Simplified schematic representation of the strategy used to generate transgenic iPSCs harbouring a DOX-inducible Tet-One system for the upregulation of the ETS-transcription factors ETV2, FLI1, and ERG. Constructs are integrated into the genome using PiggyBac transposase. (**B**) Expression analysis of nuclear immunofluorescence intensity for ERG, ETV2 and FLI1 in a selected clonal line following DOX-induction. Each dot represents the nucleus of a single cell. Bars indicate mean and standard deviation. (**C**) Stepwise DOX induction of transgenic iPSCs carrying a Tet-One inducible fluorescent reporter (exemplified by GFP) and differentiated towards iBMEC-*like* cells. Fluorescence intensity was measured daily by fluorescence microscopy.

**Fig. S2.**
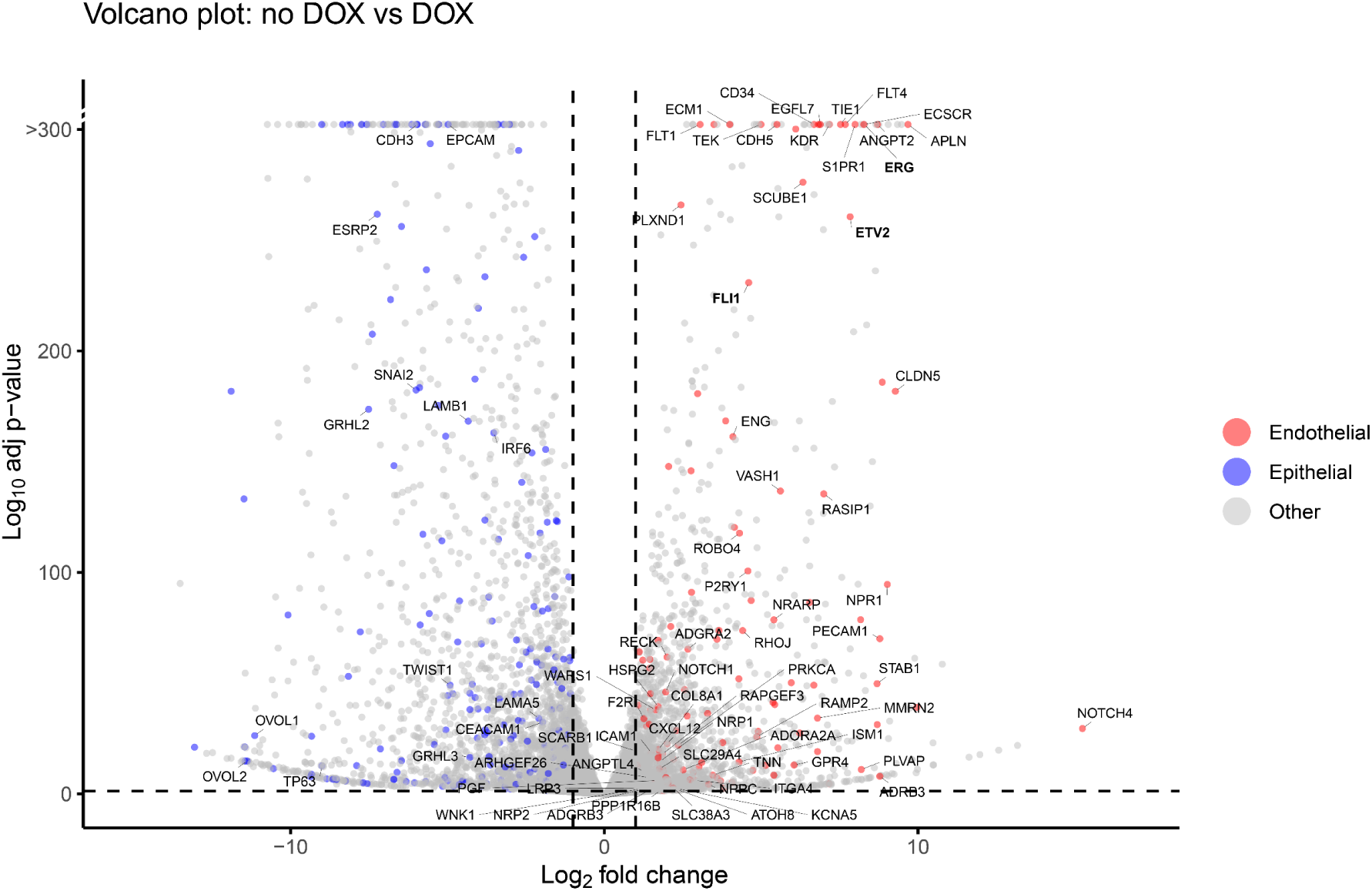
Differential gene expression between non-induced iBMEC-*like* cells and DOX-induced ETS-iBMECs. Enhanced Volcano plot showing differentially expressed genes (adjusted p-value < 0.05, Log_2_ fold change > ±1) between both cell types at day 11 of differentiation. Genes associated with endothelial (red) or epithelial (blue) GO terms are color-coded. Selected endothelial and epithelial genes are labelled with their gene symbols.

**Fig. S3.**
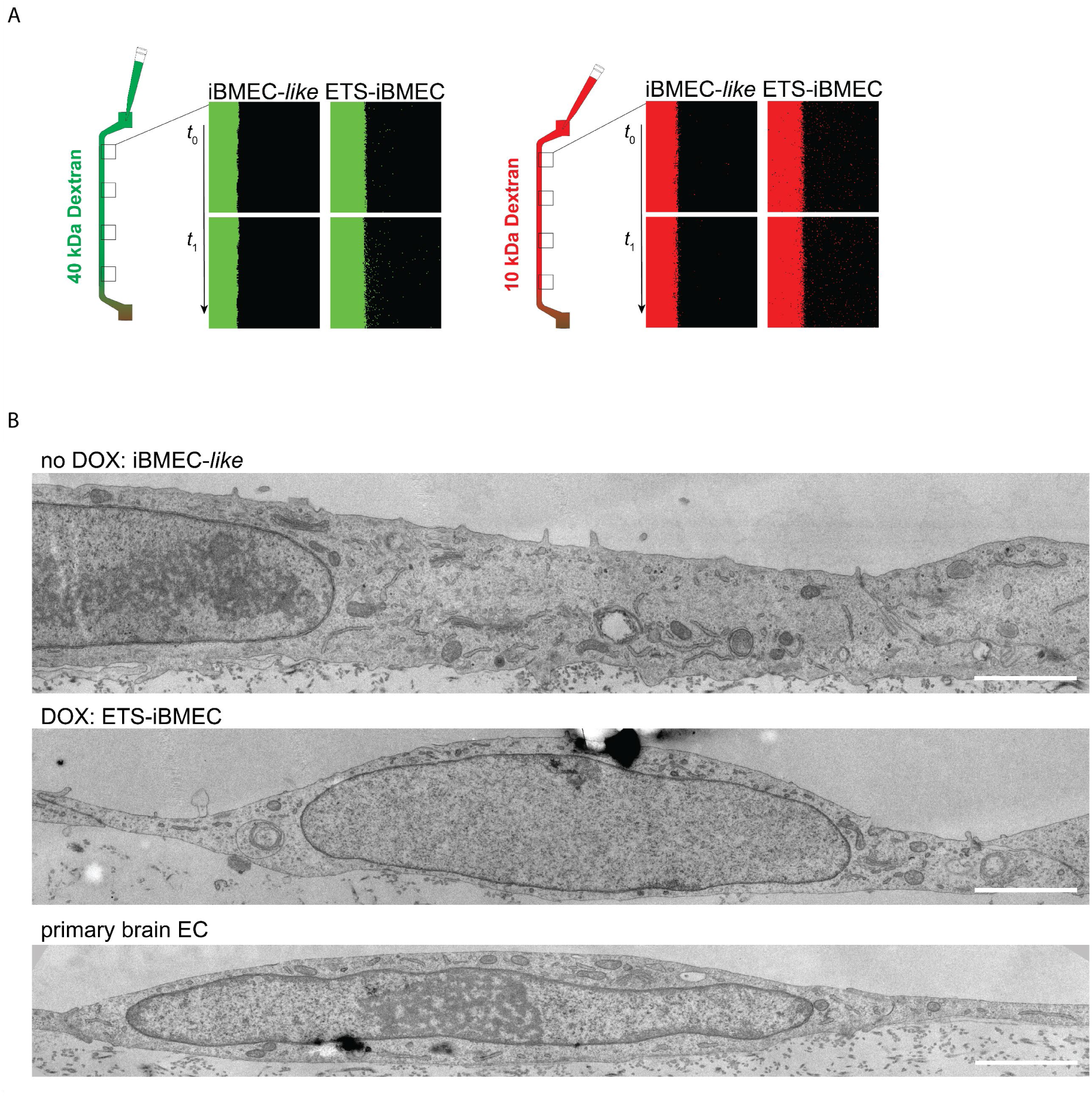
Functional and ultrastructural assays in 3D *in vitro* BBB microvessels. (**A**) Permeability of iBMEC-*like* and ETS-iBMEC microvessels perfused with fluorescently labelled 40 kDa FITC dextran (green) 10 kDa AF647 dextran (red). Apparent permeability is analysed in squared regions at two different timepoints of perfusion (t_1_ = 150 seconds). (**B**) TEM imaging of iBMEC-*like*, ETS-iBMEC and primary brain EC (scale bars: 2 µm).

**Fig. S4.**
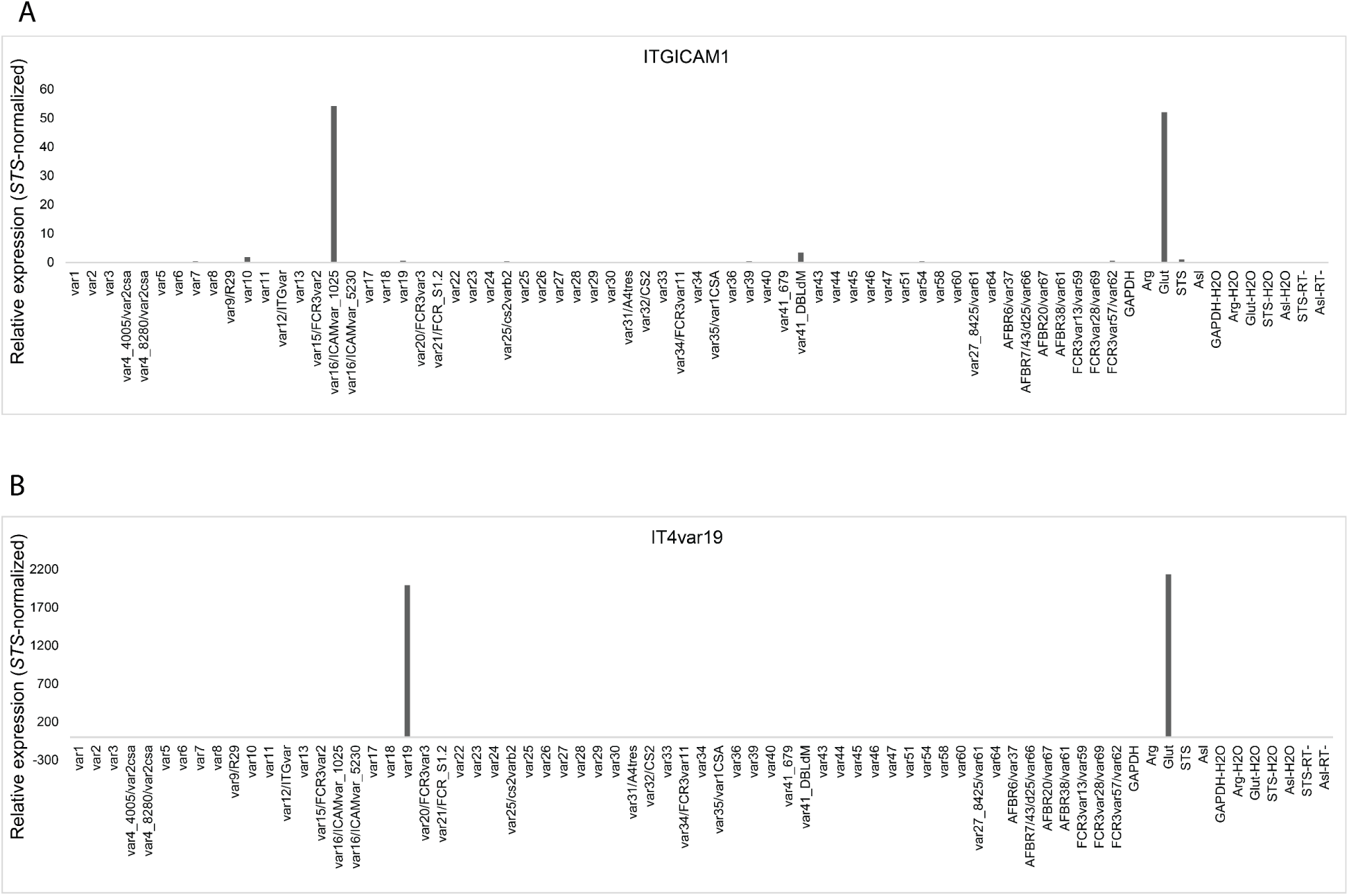
PfEMP1 variant predominant expression analysis by real-time PCR. Confirmation of (**A**) ITGICAM1 and (**B**) IT4var19 expression with *var* strain-specific primers, together with housekeeping genes to confirm the correct *var* gene expression. Relative gene expression normalized to *STS*.

**Fig. S5.**
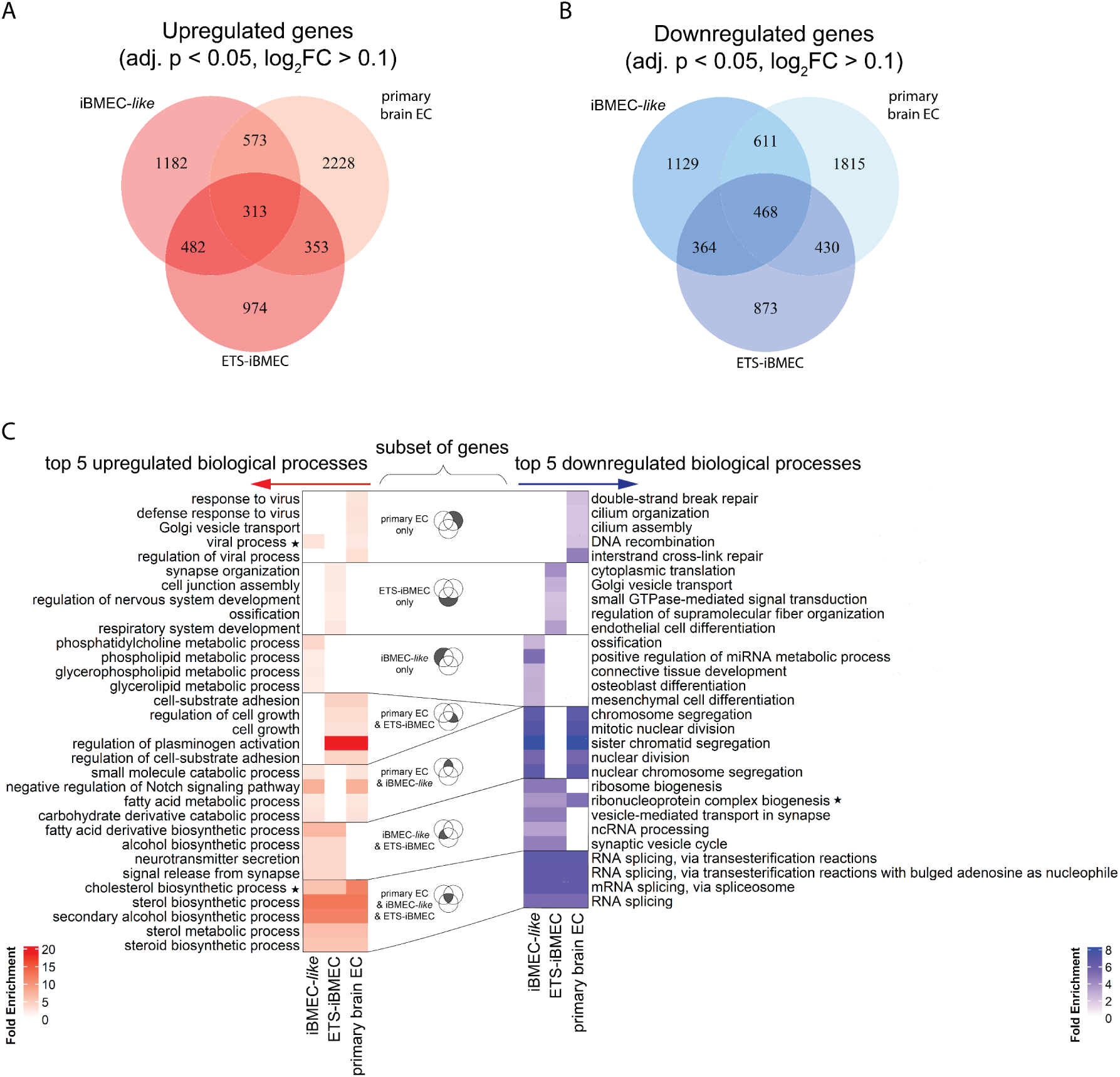
Differential gene expression in iBMEC-*like* cells, ETS-iBMEC and primary brain ECs after 24-hour malaria egress product exposure. Venn diagrams showing the overlap of significantly upregulated (**A**) and downregulated (**B**) genes (adjusted p value < 0.05, log_2_ fold change > ± 0.1) among the three cell types following exposure. (**C**) The top 5 enriched GO biological process terms for each subset of differentially expressed genes identified in the Venn diagrams. Heatmaps show fold enrichment of each GO term for the relevant cell-type-specific or shared gene subsets. Star indicates GO term expression in another Venn diagram subset.

**Fig. S6.**
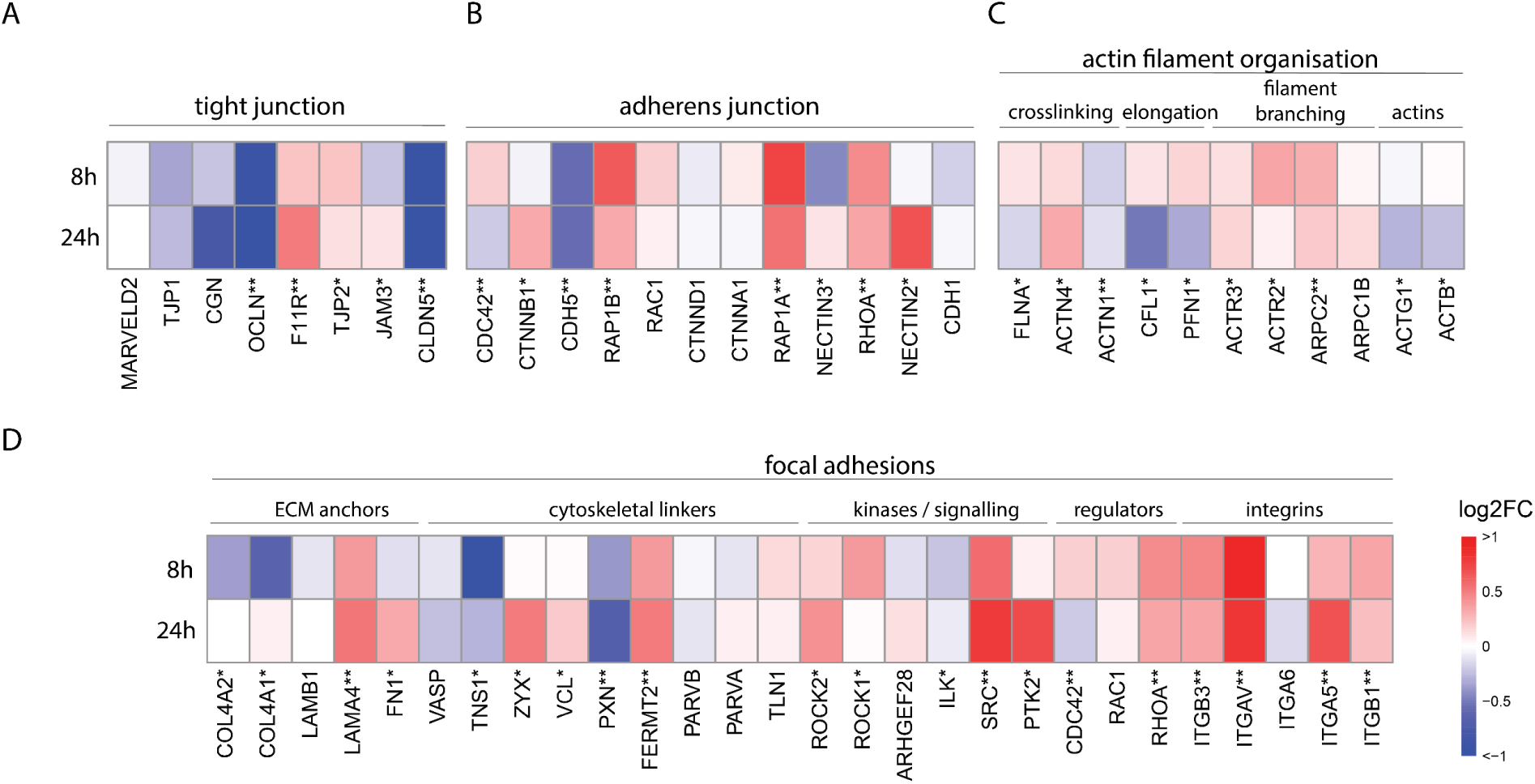
Transcriptional response of primary brain ECs to malaria egress products. Heatmaps show log_2_ fold changes in key genes related to tight junctions (**A**), adherens junctions (**B**), actin filaments (**C**), and focal adhesions (**D**) after 8- and 24-hour egress product exposure versus controls. Asterisks denote significant DESeq2 differential expression (adjust. p-value <0.05) at one (*) or both (**) time points.

## Notes

### Competing Interest Statement

The authors have declared no competing interest.

### Summary of Updates

Typo corrections only. No changes to the data or conclusions.

## Literature

1. W. H. Organization, World malaria report 2023. (World Health Organization, 2023).

2. M. F. Sabbagh, J. Nathans, A genome-wide view of the de-differentiation of central nervous system endothelial cells in culture. Elife 9, e51276 (2020).

3. E. S. Egan et al., Malaria. A forward genetic screen identifies erythrocyte CD55 as essential for Plasmodium falciparum invasion. Science 348, 711–714 (2015).

4. E. S. Egan, Beyond Hemoglobin: Screening for Malaria Host Factors. Trends Genet 34, 133–141 (2018).

5. A. Pance et al., Novel stem cell technologies are powerful tools to understand the impact of human factors on Plasmodium falciparum malaria. Front Cell Infect Microbiol 13, 1287355 (2023).

6. E. S. Lippmann et al., Derivation of blood-brain barrier endothelial cells from human pluripotent stem cells. Nature biotechnology 30, 783–791 (2012).

7. T.-E. Park et al., Hypoxia-enhanced Blood-Brain Barrier Chip recapitulates human barrier function and shuttling of drugs and antibodies. Nature communications 10, 2621 (2019).

8. T. M. Lu et al., Pluripotent stem cell-derived epithelium misidentified as brain microvascular endothelium requires ETS factors to acquire vascular fate. Proceedings of the National Academy of Sciences 118, e2016950118 (2021).

9. Y. Ding, S. P. Palecek, E. V. Shusta, iPSC-derived blood-brain barrier modeling reveals APOE isoform-dependent interactions with amyloid beta. Fluids and Barriers of the CNS 21, 79 (2024).

10. P. P. Mondkar, H. S. Seo, T. P. Lodge, S. M. Azarin, Diblock Copolymers of Poly (ethylene oxide)-b-poly (propylene oxide) Stabilize a Blood–Brain Barrier Model under Oxidative Stress. Molecular pharmaceutics 21, 5646–5660 (2024).

11. H. Nishihara et al., Intrinsic blood–brain barrier dysfunction contributes to multiple sclerosis pathogenesis. Brain 145, 4334–4348 (2022).

12. A. Gopinadhan et al., A human pluripotent stem cell-derived in vitro model of the blood-brain barrier in cerebral malaria. Fluids Barriers CNS 21, 38 (2024).

13. K. Wang et al., Robust differentiation of human pluripotent stem cells into endothelial cells via temporal modulation of ETV2 with modified mRNA. Sci Adv 6, eaba7606 (2020).

14. T. M. Lu et al., Human Induced Pluripotent Stem Cell-Derived Brain Endothelial Cells: Current Controversies. Front Physiol 12, 642812 (2021).

15. R. Morita et al., ETS transcription factor ETV2 directly converts human fibroblasts into functional endothelial cells. Proc Natl Acad Sci U S A 112, 160–165 (2015).

16. A. G. Lindgren, M. B. Veldman, S. Lin, ETV2 expression increases the efficiency of primitive endothelial cell derivation from human embryonic stem cells. Cell Regen 4, 1 (2015).

17. H. Zhang, T. Yamaguchi, Y. Kokubu, K. Kawabata, Transient ETV2 Expression Promotes the Generation of Mature Endothelial Cells from Human Pluripotent Stem Cells. Biol Pharm Bull 45, 483–490 (2022).

18. N. Koyano-Nakagawa, D. J. Garry, Etv2 as an essential regulator of mesodermal lineage development. Cardiovascular research 113, 1294–1306 (2017).

19. W. Gong et al., ETV2 functions as a pioneer factor to regulate and reprogram the endothelial lineage. Nature cell biology 24, 672–684 (2022).

20. N. Nagai et al., Downregulation of ERG and FLI1 expression in endothelial cells triggers endothelial-to-mesenchymal transition. PLoS genetics 14, e1007826 (2018).

21. D. D. Spyropoulos et al., Hemorrhage, impaired hematopoiesis, and lethality in mouse embryos carrying a targeted disruption of the Fli1 transcription factor. Molecular and cellular biology 20, 5643–5652 (2000).

22. M. J. Stebbins et al., Differentiation and characterization of human pluripotent stem cell-derived brain microvascular endothelial cells. Methods 101, 93–102 (2016).

23. G. Helms, A. K. Dasanna, U. S. Schwarz, M. Lanzer, Modeling cytoadhesion of Plasmodium falciparum-infected erythrocytes and leukocytes-common principles and distinctive features. FEBS Lett 590, 1955–1971 (2016).

24. M. Bernabeu, J. D. Smith, EPCR and Malaria Severity: The Center of a Perfect Storm. Trends Parasitol 33, 295–308 (2017).

25. Y. Zheng et al., In vitro microvessels for the study of angiogenesis and thrombosis. Proc Natl Acad Sci U S A 109, 9342–9347 (2012).

26. L. Piatti et al., Pathogenic mechanisms of Plasmodium falciparum egress unveiled by a microvascular 3D blood-brain barrier model. bioRxiv, 2024.2010.2015.618439 (2024).

27. R. K. Long et al., Plasmodium falciparum disruption of pericyte angiopoietin-1 secretion contributes to barrier breakdown in a 3D brain microvessel model. bioRxiv, 2024.2003. 2029.587334 (2024).

28. T. Lavstsen et al., Plasmodium falciparum erythrocyte membrane protein 1 domain cassettes 8 and 13 are associated with severe malaria in children. Proc Natl Acad Sci U S A 109, E1791–1800 (2012).

29. L. Turner et al., Severe malaria is associated with parasite binding to endothelial protein C receptor. Nature 498, 502–505 (2013).

30. M. Bernabeu et al., Severe adult malaria is associated with specific PfEMP1 adhesion types and high parasite biomass. Proc Natl Acad Sci U S A 113, E3270–3279 (2016).

31. A. Kessler et al., Linking EPCR-Binding PfEMP1 to Brain Swelling in Pediatric Cerebral Malaria. Cell Host Microbe 22, 601–614 e605 (2017).

32. P. K. Sahu, et al., Determinants of brain swelling in pediatric and adult cerebral malaria. JCI Insight 6, (2021).

33. M. R. Gillrie et al., Src-family kinase dependent disruption of endothelial barrier function by Plasmodium falciparum merozoite proteins. Blood 110, 3426–3435 (2007).

34. M. R. Gillrie et al., Plasmodium falciparum histones induce endothelial proinflammatory response and barrier dysfunction. Am J Pathol 180, 1028–1039 (2012).

35. J. Gallego-Delgado et al., Angiotensin receptors and beta-catenin regulate brain endothelial integrity in malaria. J Clin Invest 126, 4016–4029 (2016).

36. J. Tapial et al., An atlas of alternative splicing profiles and functional associations reveals new regulatory programs and genes that simultaneously express multiple major isoforms. Genome Res 27, 1759–1768 (2017).

37. K. B. Seydel et al., Brain swelling and death in children with cerebral malaria. N Engl J Med 372, 1126–1137 (2015).

38. P. K. Sahu et al., Brain magnetic resonance imaging reveals different courses of disease in pediatric and adult cerebral malaria. Clinical Infectious Diseases 73, e2387–e2396 (2021).

39. S. Mohanty et al., Evidence of Brain Alterations in Noncerebral Falciparum Malaria. Clin Infect Dis 75, 11–18 (2022).

40. C. Gomes, et al., Endothelial transcriptomic analysis identifies biomarkers of severe and cerebral malaria. JCI Insight 8, (2023).

41. A. K. Tripathi, W. Sha, V. Shulaev, M. F. Stins, D. J. Sullivan, Jr., Plasmodium falciparum-infected erythrocytes induce NF-kappaB regulated inflammatory pathways in human cerebral endothelium. Blood 114, 4243–4252 (2009).

42. M. Zuniga et al., Plasmodium falciparum and TNF-alpha Differentially Regulate Inflammatory and Barrier Integrity Pathways in Human Brain Endothelial Cells. mBio 13, e0174622 (2022).

43. C. Howard, F. Joof, R. Hu, J. D. Smith, Y. Zheng, Probing cerebral malaria inflammation in 3D human brain microvessels. Cell Rep 42, 113253 (2023).

44. J. M. Gomez-Salinero et al., Cooperative ETS Transcription Factors Enforce Adult Endothelial Cell Fate and Cardiovascular Homeostasis. Nat Cardiovasc Res 1, 882–899 (2022).

45. Y. Ding et al., ETV2 Overexpression Promotes Efficient Differentiation of Pluripotent Stem Cells to Endothelial Cells. Biotechnol Bioeng 122, 1914–1928 (2025).

46. D. Chen et al., Pioneer factor ETV2 safeguards endothelial cell specification by recruiting the repressor REST to restrict alternative lineage commitment. Nat Cardiovasc Res 4, 689–709 (2025).

47. L. Gong et al., Rapid generation of functional vascular organoids via simultaneous transcription factor activation of endothelial and mural lineages. Cell Stem Cell, (2025).

48. A. H. M. Ng et al., A comprehensive library of human transcription factors for cell fate engineering. Nat Biotechnol 39, 510–519 (2021).

49. J. M. Gomez-Salinero, D. Redmond, S. Rafii, Microenvironmental determinants of endothelial cell heterogeneity. Nat Rev Mol Cell Biol 26, 476–495 (2025).

50. S. Rieck et al., Forward programming of human induced pluripotent stem cells via the ETS variant transcription factor 2: rapid, reproducible, and cost-effective generation of highly enriched, functional endothelial cells. Cardiovasc Res 120, 1472–1484 (2024).

51. A. C. Luo et al., A streamlined method to generate endothelial cells from human pluripotent stem cells via transient doxycycline-inducible ETV2 activation. Angiogenesis 27, 779–795 (2024).

52. A. Cui et al., Generation of hiPSC-derived brain microvascular endothelial cells using a combination of directed differentiation and transcriptional reprogramming strategies. bioRxiv, (2024).

53. J. Storm et al., Cerebral malaria is associated with differential cytoadherence to brain endothelial cells. EMBO Mol Med 11, (2019).

54. M. J. Ponsford et al., Sequestration and microvascular congestion are associated with coma in human cerebral malaria. J Infect Dis 205, 663–671 (2012).

55. J. Hanson et al., Microvascular obstruction and endothelial activation are independently associated with the clinical manifestations of severe falciparum malaria in adults: an observational study. BMC Med 13, 122 (2015).

56. A. M. Dondorp et al., Direct in vivo assessment of microcirculatory dysfunction in severe falciparum malaria. J Infect Dis 197, 79–84 (2008).

57. J. Hanson et al., Relative contributions of macrovascular and microvascular dysfunction to disease severity in falciparum malaria. J Infect Dis 206, 571–579 (2012).

58. M. Avril, M. Bernabeu, M. Benjamin, A. J. Brazier, J. D. Smith, Interaction between Endothelial Protein C Receptor and Intercellular Adhesion Molecule 1 to Mediate Binding of Plasmodium falciparum-Infected Erythrocytes to Endothelial Cells. mBio 7, (2016).

59. A. R. Berendt, D. L. Simmons, J. Tansey, C. I. Newbold, K. Marsh, Intercellular adhesion molecule-1 is an endothelial cell adhesion receptor for Plasmodium falciparum. Nature 341, 57–59 (1989).

60. J. A. Rowe, A. Claessens, R. A. Corrigan, M. Arman, Adhesion of Plasmodium falciparum-infected erythrocytes to human cells: molecular mechanisms and therapeutic implications. Expert Rev Mol Med 11, e16 (2009).

61. M. Bernabeu et al., Binding Heterogeneity of Plasmodium falciparum to Engineered 3D Brain Microvessels Is Mediated by EPCR and ICAM-1. mBio 10, (2019).

62. R. A. Reyes et al., Broadly inhibitory antibodies to severe malaria virulence proteins. Nature 636, 182–189 (2024).

63. F. Joof et al., Plasma From Older Children in Malawi Inhibits Plasmodium falciparum Binding in 3-Dimensional Brain Microvessels. J Infect Dis 230, e1402–e1411 (2024).

64. K. Dorovini-Zis et al., The Neuropathology of Fatal Cerebral Malaria in Malawian Children. The American Journal of Pathology 178, 2146–2158 (2011).

65. H. Brown et al., Blood-brain barrier function in cerebral malaria in Malawian children. Am J Trop Med Hyg 64, 207–213 (2001).

66. H. Brown et al., Evidence of blood-brain barrier dysfunction in human cerebral malaria. Neuropathol Appl Neurobiol 25, 331–340 (1999).

67. T. Walchli et al., Single-cell atlas of the human brain vasculature across development, adulthood and disease. Nature 632, 603–613 (2024).

68. V. Jain et al., Plasma IP-10, apoptotic and angiogenic factors associated with fatal cerebral malaria in India. Malar J 7, 83 (2008).

69. A. L. Conroy et al., Whole blood angiopoietin-1 and -2 levels discriminate cerebral and severe (non-cerebral) malaria from uncomplicated malaria. Malar J 8, 295 (2009).

70. A. L. Conroy et al., Angiopoietin-2 levels are associated with retinopathy and predict mortality in Malawian children with cerebral malaria: a retrospective case-control study*. Crit Care Med 40, 952–959 (2012).

71. F. E. Lovegrove et al., Serum angiopoietin-1 and -2 levels discriminate cerebral malaria from uncomplicated malaria and predict clinical outcome in African children. PLoS One 4, e4912 (2009).

72. T. W. Yeo et al., Angiopoietin-2 is associated with decreased endothelial nitric oxide and poor clinical outcome in severe falciparum malaria. Proc Natl Acad Sci U S A 105, 17097–17102 (2008).

73. S. J. Higgins et al., Dysregulation of angiopoietin-1 plays a mechanistic role in the pathogenesis of cerebral malaria. Sci Transl Med 8, 358ra128 (2016).

74. T. Furuta, M. Kimura, N. Watanabe, Elevated levels of vascular endothelial growth factor (VEGF) and soluble vascular endothelial growth factor receptor (VEGFR)-2 in human malaria. Am J Trop Med Hyg 82, 136–139 (2010).

75. I. M. Medana et al., Induction of the vascular endothelial growth factor pathway in the brain of adults with fatal falciparum malaria is a non-specific response to severe disease. Histopathology 57, 282–294 (2010).

76. R. Webster et al., Adjunctive ruxolitinib attenuates inflammation and enhances antiparasitic immunity in human volunteers experimentally infected with Plasmodium falciparum. MedRxiv, 2025.2003. 2026.25324737 (2025).

77. L. Bukali et al., Transient JAK/STAT inhibition by ruxolitinib modulates malaria-specific CD4+ T cell responses and enhances recall immunity in volunteers experimentally infected with Plasmodium falciparum. medRxiv, 2025.2004. 2009.25325416 (2025).

78. M. F. Chughlay et al., Safety, Tolerability, Pharmacokinetics, and Pharmacodynamics of Coadministered Ruxolitinib and Artemether-Lumefantrine in Healthy Adults. Antimicrobial Agents and Chemotherapy 66, e01584–01521 (2022).

79. V. Barrera et al., Comparison of CD8(+) T Cell Accumulation in the Brain During Human and Murine Cerebral Malaria. Front Immunol 10, 1747 (2019).

80. B. A. Riggle et al., CD8+ T cells target cerebrovasculature in children with cerebral malaria. J Clin Invest 130, 1128–1138 (2020).

81. F. Korbmacher, M. Bernabeu, Induced pluripotent stem cell-based tissue models to study malaria: a new player in the research game. Curr Opin Microbiol 84, 102585 (2025).

82. K. Woltjen et al., piggyBac transposition reprograms fibroblasts to induced pluripotent stem cells. Nature 458, 766–770 (2009).

83. M. Avril et al., A restricted subset of var genes mediates adherence of Plasmodium falciparum-infected erythrocytes to brain endothelial cells. Proc Natl Acad Sci U S A 109, E1782–1790 (2012).

84. C. F. Ockenhouse et al., Human vascular endothelial cell adhesion receptors for Plasmodium falciparum-infected erythrocytes: roles for endothelial leukocyte adhesion molecule 1 and vascular cell adhesion molecule 1. J Exp Med 176, 1183–1189 (1992).

85. L. Piatti, C. C. Howard, Y. Zheng, M. Bernabeu, Binding of Plasmodium falciparum-Infected Red Blood Cells to Engineered 3D Microvessels. Methods Mol Biol 2470, 557–585 (2022).

86. J. H. Janes et al., Investigating the host binding signature on the Plasmodium falciparum PfEMP1 protein family. PLoS Pathog 7, e1002032 (2011).

87. R. Patro, G. Duggal, M. I. Love, R. A. Irizarry, C. Kingsford, Salmon provides fast and bias-aware quantification of transcript expression. Nat Methods 14, 417–419 (2017).

88. C. Soneson, M. I. Love, M. D. Robinson, Differential analyses for RNA-seq: transcript-level estimates improve gene-level inferences. F1000Res 4, 1521 (2015).

89. M. I. Love, W. Huber, S. Anders, Moderated estimation of fold change and dispersion for RNA-seq data with DESeq2. Genome Biol 15, 550 (2014).

90. A. Zhu, J. G. Ibrahim, M. I. Love, Heavy-tailed prior distributions for sequence count data: removing the noise and preserving large differences. Bioinformatics 35, 2084–2092 (2019).

91. R. Kolde. (2019).

92. T. Wu, et al., clusterProfiler 4.0: A universal enrichment tool for interpreting omics data. The innovation 2, (2021).

93. G. Yu. (2024).

94. K. Blighe, S. Rana, M. Lewis. (2024).

